# Subunit Communication within Dimeric SF1 DNA Helicases

**DOI:** 10.1101/2024.02.20.581227

**Authors:** Binh Nguyen, John Hsieh, Christopher J. Fischer, Timothy M. Lohman

**Affiliations:** Department of Biochemistry & Molecular Biophysics Washington University School of Medicine, 660 S. Euclid Ave., Saint Louis, MO 63110; Biochemistry & Biophysics, Blueprint Medicines, Cambridge, MA 02139; Department of Physics and Astronomy University of Kansas, Lawrence KS 66045

**Keywords:** Helicase activation, gain of function, single molecule fluorescence, Rep, UvrD

## Abstract

Monomers of the Superfamily (SF) 1 helicases, *E. coli* Rep and UvrD, can translocate directionally along single stranded (ss) DNA, but must be activated to function as helicases. In the absence of accessory factors, helicase activity requires Rep and UvrD homo-dimerization. The ssDNA binding sites of SF1 helicases contain a conserved aromatic amino acid (Trp250 in Rep and Trp256 in UvrD) that stacks with the DNA bases. Here we show that mutation of this Trp to Ala eliminates helicase activity in both Rep and UvrD. Rep(W250A) and UvrD(W256A) can still dimerize, bind DNA, and monomers still retain ATP-dependent ssDNA translocase activity, although with lower processivity than wild type monomers. Although neither wtRep monomers nor Rep(W250A) monomers possess helicase activity by themselves, using both ensemble and single molecule methods, we show that helicase activity is achieved upon formation of a Rep(W250A)/wtRep hetero-dimer. An ATPase deficient Rep monomer is unable to activate a wtRep monomer indicating that ATPase activity is needed in both subunits of the Rep hetero-dimer. We find the same results with *E. coli* UvrD and its equivalent mutant (UvrD(W256A)). Importantly, Rep(W250A) is unable to activate a wtUvrD monomer and UvrD(W256A) is unable to activate a wtRep monomer indicating that specific dimer interactions are required for helicase activity. We also demonstrate subunit communication within the dimer by virtue of Trp fluorescence signals that only are present within the Rep dimer, but not the monomers. These results bear on proposed subunit switching mechanisms for dimeric helicase activity.

## Introduction

DNA helicases/translocases are ubiquitous ATP-dependent enzymes that play essential roles in all aspects of genome maintenance, including DNA replication, recombination, and DNA repair [1-3]. These enzymes are diverse in structure, oligomeric state and activity. These include the ring-shaped hexameric helicases that encircle the DNA and function as the primary helicase at DNA replication forks [4]. However, the majority of DNA helicases are members of the non-hexameric superfamilies (SF) 1 and 2 [1]. Some SF1 and SF2 helicases, such as phage T4 Dda [5, 6] can function as monomers *in vitro*. However, the SF1 DNA helicases, *E. coli* Rep, *E. coli* UvrD, and *B. stearothermophilus* PcrA do not possess processive DNA helicase activity as monomers. Rep, UvrD and PcrA monomers can translocate directionally (3’ to 5’) along single stranded (ss) DNA while hydrolyzing ATP [7-11], yet, in order to unwind duplex DNA, these enzymes need to be activated. Such activation can occur by a number of mechanisms. Activation mechanisms *in vitro* include homo-dimerization [12-30], removal of its auto-inhibitory 2B subdomain [29, 31, 32], crosslinking the monomer into a closed conformation [33], and interaction with accessory factors PriC for Rep [29, 34], RepD for PcrA [35] and MutL for UvrD [36-40]. However, the molecular bases for enzyme activation remains unknown. For that matter, the molecular mechanism for DNA unwinding by these SF1 helicases remains unclear.

Movement of the auto-inhibitory 2B sub-domain occurs during activation of both Rep and UvrD dimers. The 2B sub-domain of Rep [41], PcrA [42, 43], and UvrD [28, 44, 45] can populate a range of rotational conformations by rotating about a hinge region connected to the 2A sub-domain by as much as from 130° to 160°, with limiting structures referred to as “open” and “closed” forms. In the case of Rep, the 2B sub-domain inhibits Rep monomer helicase activity, since removal of the 2B sub-domain activates monomer helicase activity [11, 31, 32]. The presumption is that the 2B sub-domains of UvrD and PcrA also inhibit the monomer helicase activity. In the case of *E. coli* UvrD, activation by dimerization and MutL is correlated with movement of its 2B sub-domain into a conformation that is intermediate between a fully open and fully closed conformation [28, 40]. It has been postulated that this movement through dimerization relieves the auto-inhibitory effect of the 2B sub-domain. Recent studies of the homologous SF1 UvrD1 helicase from *Mycobacterium tuberculosis* show that it also requires dimerization for helicase activity and that dimerization occurs via a redox dependent covalent Cystine crosslink between the 2B sub-domains of each monomer [46, 47].

Whereas *E. coli* UvrD can form dimers and tetramers in the absence of DNA [26]. Rep dimerization requires DNA binding [12-14] and both DNA binding and dimerization dramatically affect the ATPase activities of Rep [17-19, 48]. This indicates that the subunits within the Rep dimer communicate allosterically and this communication plays a role in regulating helicase activity.

Crystal structures of Rep [41], UvrD [44, 45], and PcrA monomers bound to DNA show a common characteristic within their ssDNA binding sites (**Figure 1**). A conserved Trp residue in motif III in the 1A sub-domain (RepW250 and UvrDW256 and PcrAW259) interacts directly with DNA via a stacking interaction with the nucleotide bases [41, 43, 44] as shown in **Figure 1**. This Trp residue has been proposed to play a critical role in DNA translocation/unwinding [43, 49, 50]. Mutation of this Trp to Ala in PcrA (W259A) results in defects in DNA binding [50], although the helicase and ssDNA translocase activities of this mutant were not examined.

**Figure 1.**
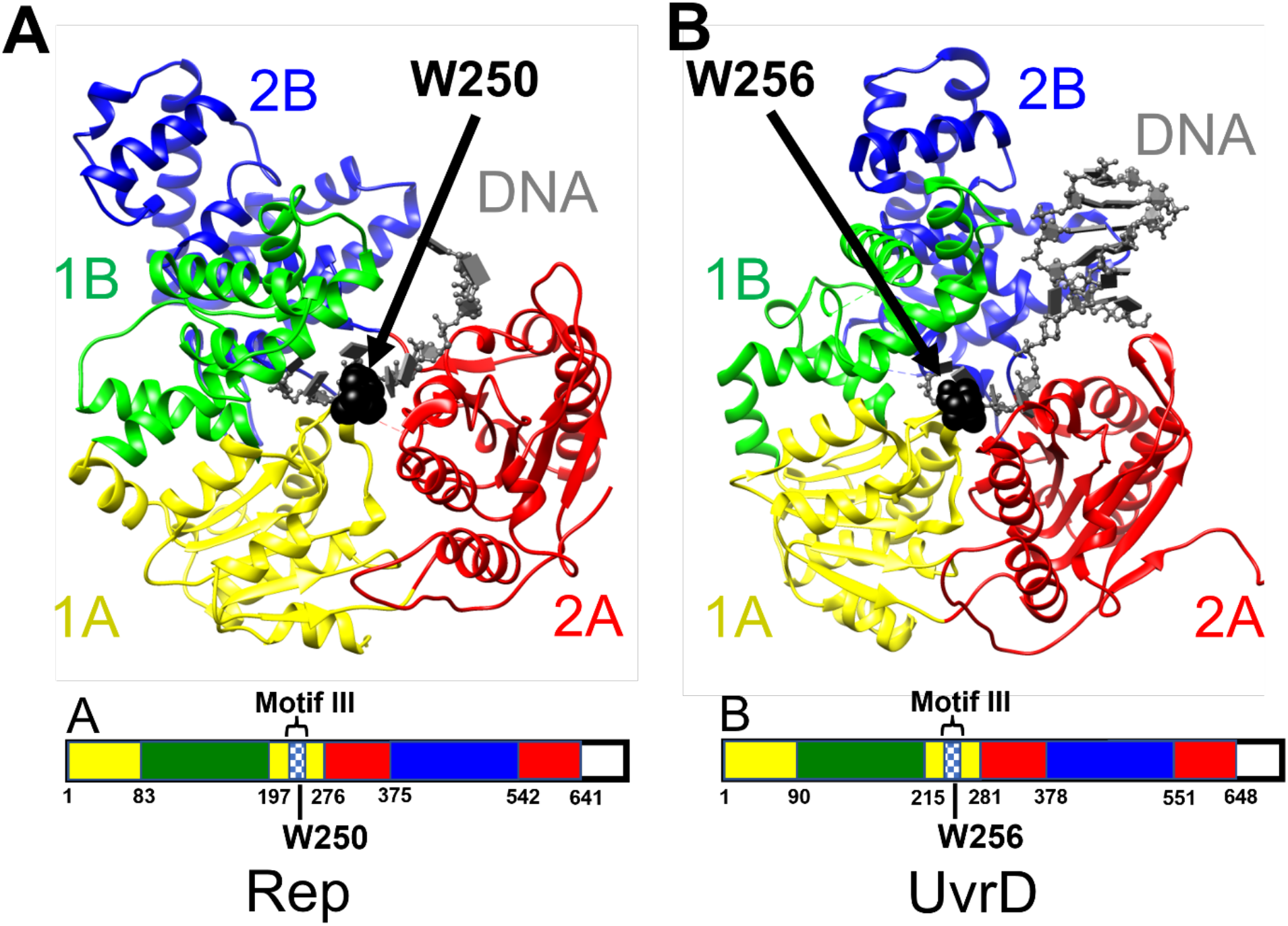
The conserved Trp within motif III in Rep and UvrD interacts with ssDNA. Crystal structures of (**A**) Rep (1UAA) and (**B**) UvrD (2IS6) with the four sub-domains colored in yellow (1A), red (2A), green (1B) and blue (2B). The conserved Trp (RepW250 and UvrDW256 in black) located within motif III in the 1A sub-domain forms a stacking interaction with ssDNA.

In this study we have mutated this conserved Trp to Ala in both Rep and UvrD and have characterized their DNA binding and enzymatic activities. Rep(W250A) and UvrD(W256A) are completely defective for DNA helicase activity. The mutants possess weaker DNA binding and ATPase activity, but still undergo dimerization, yet the dimers are inactive as helicases. Rep(W250A) and UvrD(W256A) monomers are able to undergo directional (3’ to 5’) ATP-dependent ssDNA translocation, although with lower processivity than the wt monomers. Most interestingly, Rep(W250A) can form a hetero-dimer with a wtRep monomer which activates the DNA helicase activity. Hetero-dimerization of UvrD(W256A) with wtUvrD monomer also activates helicase activity. Cross activation of wtUvrD monomer by Rep(W250A) or of wtRep monomer by UvrD(W256A) does not occur indicating that specific interactions and communication between subunits within each dimer are needed for activation.

## Results

### Trp-250 of Rep is responsible for Trp fluorescence changes upon DNA binding

Upon binding ssDNA, Rep’s Trp fluorescence is quenched by ∼20% [51]. Wild type Rep contains eight Trp residues, however, based on crystal structures of Rep bound to ssDNA, only Trp250 interacts directly with ssDNA via stacking interaction as shown in **Figure 1A** [41]. To determine if Trp250 is responsible for this fluorescence quenching we mutated Trp250 to Ala and examined the binding of Rep(W250A) to ssDNA. **Figure 2A** shows stopped-flow kinetic traces indicating that no Trp fluorescence change is observed upon mixing Rep(W250A) with a 16-nucleotide ssDNA (5’-(dT)_5_-(2-AP)-(dT)_4_-(2-AP)- (dT)_5_), whereas Trp fluorescence is quenched for wtRep binding to the DNA.

**Figure 2.**
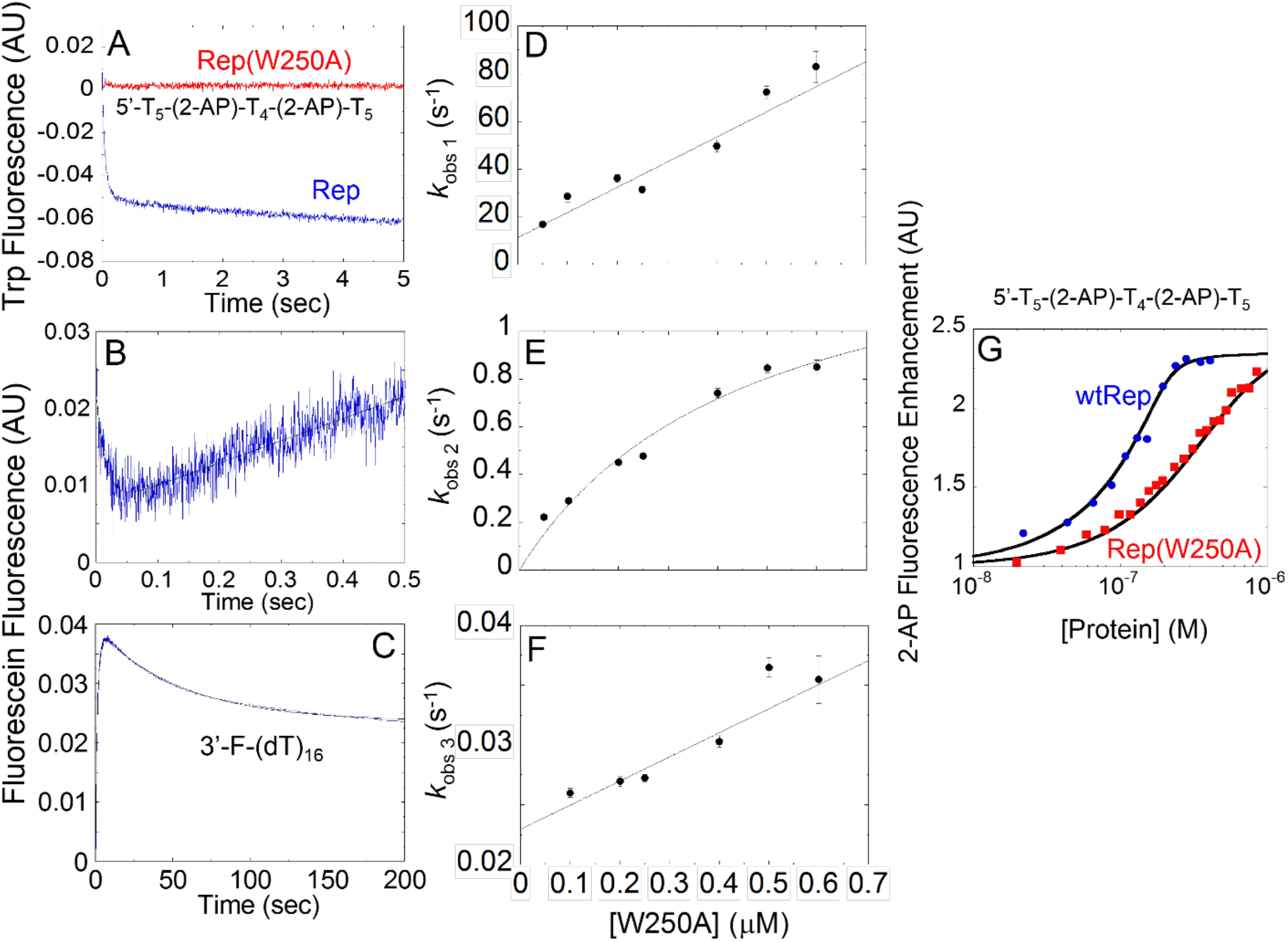
Rep(W250A) can bind ssDNA and dimerize. (**A**)-Stopped-flow mixing experiments show that RepW250 is responsible for the Trp fluorescence change upon Rep (125 nM) binding to 5’-(dT)_5_-(2-AP)-(dT)_4_-(2-AP)-(dT)_5_ (600 nM) (buffer A, 4 ºC). (**B)** and (**C**)-Binding of Rep(W250A) (400 nM) to 3’-Fluorescein-labeled ssDNA (3’-F-(dT)_16_, 10 nM) monitored by the change in fluorescein fluorescence (λ_ex_ = 492 nm, λ_em_ ≥ 520 nm) (buffer A 4 ºC) shows three phases representing binding, a conformational change and dimerization. (**D**), (**E**) and (**F**)- The three observed rate constants plotted as a function of [Rep(W250A)]. (**D**)- linear dependence of *k*_obs,1_ on [Rep(W250A)] with slope indicating *k*_1_ = 8.7±0.6 ×10^7^ M^-1^ s^-1^, (**E**)- hyperbolic dependence of *k*_obs,2_ on [Rep(W250A)]. (**F**)-linear dependence of *k*_obs,3_ on [Rep(W250A)] indicating a dimerization rate constant of 2.7±0.2 ×10^4^ M^-1^ s^-1^. (**G**) Equilibrium fluorescence titrations of (5’-(dT)_5_-(2-AP)-(dT)_4_-(2-AP)-(dT)_5_) (100 nM) with Rep(W250A) and wtRep (buffer A, 4 ºC). The solid lines are simulations based on the equilibrium constants in **Scheme S1 (Table S2)**.

The absence of Trp fluorescence quenching for Rep(W250A) is not due to an inability to bind ssDNA or to dimerize. We used a stopped-flow kinetic approach to examine binding of Rep(W250A) to a fluorescein labeled ssDNA (3’-F-(dT)_16_) and its subsequent dimerization [51]. As shown in **Figure 2B,C**, Rep(W250A) binding to 3’-F- (dT)_16_ results in a triphasic fluorescein fluorescence change, similar to what was observed with wtRep [51]. An initial decrease in fluorescein fluorescence is followed by a transient increase and then a final fluorescence decrease. As shown previously [51], the first phase represents the initial binding of Rep(W250A) to 3’-F-(dT)_16_ to form a PS monomer complex. The second phase reflects an isomerization event (PS to PS*), and the third phase reflects ssDNA binding-induced dimerization to form a P_2_S complex. Experiments were performed at a constant 3’-F-(dT)_16_ concentration (10 nM) with increasing concentrations of Rep(W250A) and the three relaxation times (k_obs,1_, k_obs,2_, k_obs,3_) are plotted vs. [Rep(W250A)] in **Figure 2D-F**. These data were analyzed using **Scheme S1** to obtain the individual rate constants (Supplemental **Table S1**). When compared to previous studies of wtRep [51], the major differences are in the monomer dissociation rate constants, k_-1_, which is ∼10-fold higher for Rep(W250A) and in the rate constant for dimer formation, k_3_, which is ∼10 fold lower for Rep(W250A).

Additional kinetic and equilibrium studies were performed with (5’-(dT)_5_-(2-AP)- (dT)_4_-(2-AP)-(dT)_5_) (Supplemental **Figures S1** and **S2**) to resolve the isomerization constants (*k*_*2*_ and *k*_*-2*_). This DNA contains the fluorescent base 2 aminopurine (2-AP), which undergoes a fluorescence enhancement upon binding Rep [51]. Experiments in Supplemental **Figure S1** were performed by mixing a constant amount of Rep(W250A) (100 nM) with an increasing concentration of ssDNA to minimize Rep(W250A) dimerization on DNA. The rate constants (*k*_*1*_ and *k*_*-1*_) for the binding of Rep(W250A) to DNA were determined as shown in Supplemental **Figure S1**. Experiments in Supplemental **Figure S2** were performed by mixing a constant of amount of ssDNA (100 nM) with increasing Rep(W250A) concentrations, which provide data on the rate of Rep(W250A) dimerization. Global fitting of the kinetic traces in **Figures S1** and **S2** to **Scheme S1** yield the rate constants and equilibrium constants (K_1_, K_2_ and K_3_) reported in **Table S2. Figure 2G** shows the results of equilibrium titrations of 5’-(dT)_5_-(2-AP)-(dT)_4_- (2-AP)-(dT)_5_ with wtRep and Rep(W250A) monitoring 2-AP fluorescence enhancement, showing weaker overall binding of Rep(W250A). These equilibrium titrations are well described by **Scheme S1** and the three equilibrium constants (K_1_, K_2_, K_3_) obtained from the kinetics experiments. The reduction in the overall binding affinity of the Rep(W250A) compared to the wtRep results from a reduction in all three equilibrium constants. However, Rep(W250A) is able to bind ssDNA and dimerize although no Trp fluorescence quenching accompanies binding and dimerization. Hence, Trp250 is solely responsible for the Trp fluorescence quenching that occurs upon wtRep binding to ssDNA.

### Mutations of the conserved Trp in the DNA binding sites of Rep and UvrD abolish helicase activity

We examined the helicase activity of Rep(W250A) using a fluorescence stopped-flow DNA unwinding assay (**Figure 3**). An 18-bp duplex DNA possessing a 3’- (dT)_20_ single stranded flanking region with a Cy5 and a BHQ2 quencher at the blunt end (3’-(dT)_20_-18bp Cy5/BHQ2, see Methods) was used as the DNA unwinding substrate. The Cy5 fluorescence is quenched when DNA is in the duplex form. Upon DNA unwinding, the Cy5-labeled strand is separated from the quencher labeled strand resulting in Cy5 fluorescence enhancement (**Figure 3A**). Protein (800 nM) was premixed with the DNA (200 nM) in one syringe. The mixture was rapidly mixed with ATP and protein trap (3’- (dT)_40_-HP10, see Methods section) in the other syringe. The protein trap prevents rebinding of protein to the DNA substrate, ensuring single round unwinding kinetics. **Figure 3B** shows the expected DNA unwinding kinetics when wtRep is in excess over the DNA substrate, whereas Rep(W250A) displays no helicase activity even when in large excess. Similarly, no DNA unwinding was observed with a large excess of UvrD(W256A) over the DNA substrate (**Figure 3C**). Therefore, mutation of this Trp residue to Ala abolishes helicase activity in both Rep and UvrD.

**Figure 3.**
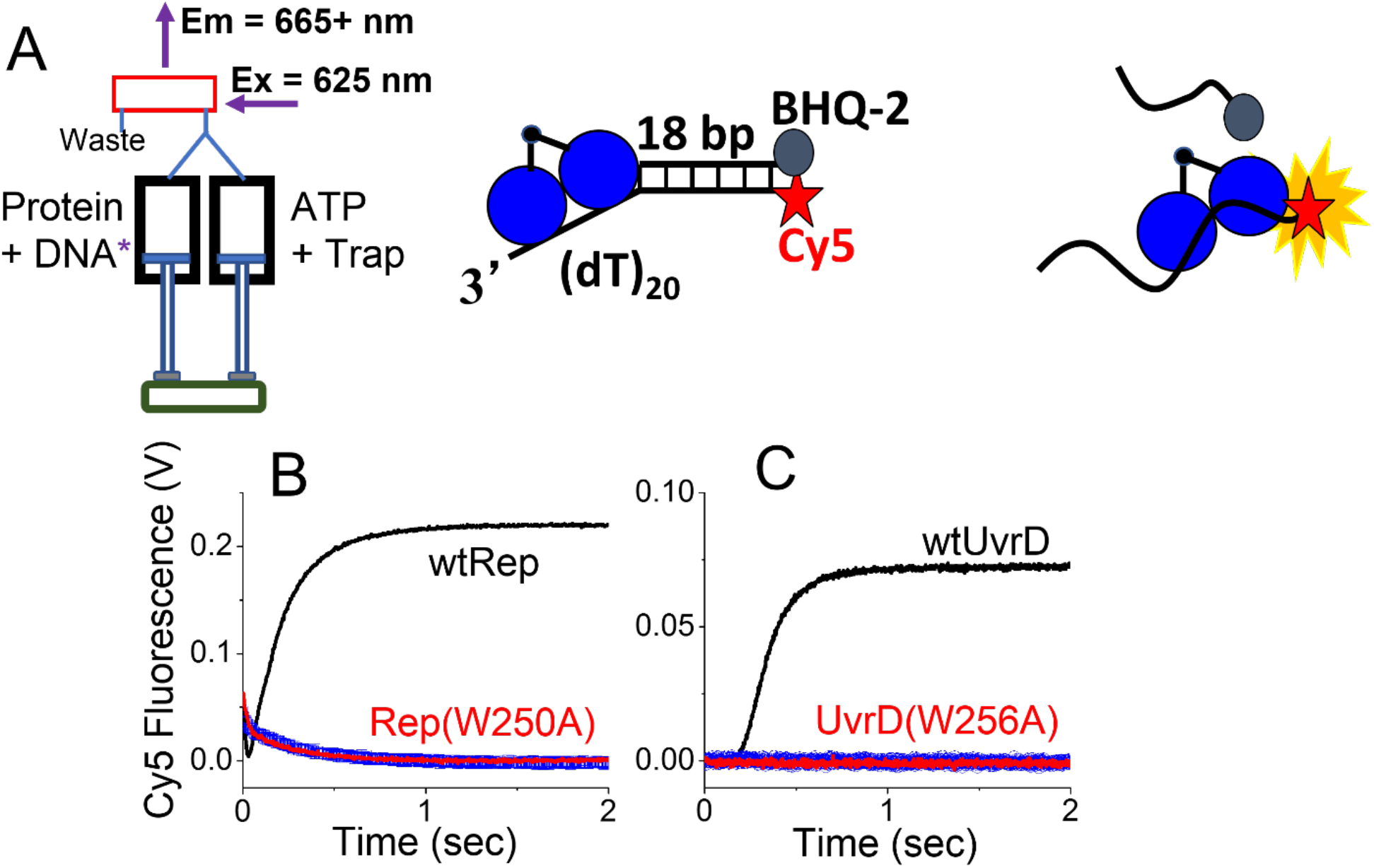
Rep(W250A) and UvrD(W256A) do not possess helicase activity. (**A**) Schematic of the fluorescence stopped-flow DNA unwinding experiment using a DNA substrate (3’-(dT)_20_-18bp Cy5/BHQ2) in which the ends of the DNA are labeled with Cy5 and a black hole quencher (BHQ2). DNA unwinding and strand separation is accompanied by an increase in Cy5 fluorescence. (**B**) DNA unwinding time courses for DNA (100 nM) with wtRep (400 nM) and Rep(W250A) (400 nM). (**C**) DNA unwinding time courses for DNA (100 nM) with wtUvrD (400 nM) and UvrD(W256A) (400 nM) in the presence of 5 μM 3’-(dT)_40_-HP10 as a protein trap (Buffer U, 25 ºC).

*E. coli* Rep helicase is essential for replication of bacteriophage φX174 [52]. Thus, we also examined the ability of Rep(W250A) to support φX174 phage replication using a φX174 phage plaque-forming assay [53]. An *E. coli* strain, *CK11Δrep/pIWcI*, containing a deletion of the *rep* gene [54] was transformed with plasmids pRepO, expressing wtRep [55] or pJH1, expressing Rep(W250A) [54]. Successful φX174 replication in the *CK11Δrep/pIWcI* results in the lysis of *E. coli*. The results in **Table 1** indicate that φX174 phage was not able to replicate in the *rep* deletion strain *CK11Δrep/pIWcI* as expected. *CK11Δrep/pIWcI* transformed with pRepO, expressing wtRep, supported phage replication (70 plaques). However, *CK11Δrep/pIWcI* transformed with pJH1, expressing Rep(W250A) showed no plaques indicating that Rep(W250A) also shows no helicase activity *in vivo*. Substitution of Trp250 with Phe also supports φX174 phage replication (**Table 1**) suggesting the importance of an aromatic residue at that position.

**Table 1.**
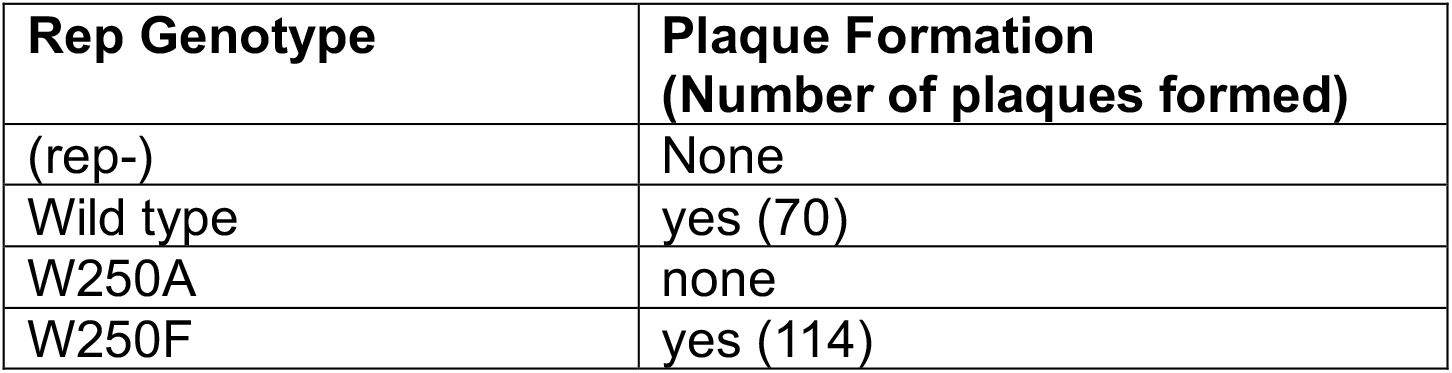
W250A mutation does not support phage replication.

| **Rep Genotype** | **Plaque Formation**<br>**(Number of plaques formed)** |
| --- | --- |
| (rep-) | None |
| Wild type | yes (70) |
| W250A | none |
| W250F | yes (114) |

### Rep(W250A) can bind and hydrolyze ATP

The steady state ATPase activity of wtRep is strongly dependent on DNA binding and Rep dimerization [19]. We therefore examined the ssDNA-dependent ATPase activities of both the Rep(W250A) monomer (PS/PS*) and the dimer (P_2_S) as described previously for wtRep [19]. The PS complex was formed by mixing low concentrations of Rep(W250A) (5-50 nM) with a large excess of (dT)_16_ (4 μM). The P_2_S complex was formed by mixing a large excess of Rep(W250A) (800 nM) with low concentrations of (dT)_16_ (1 -10 nM). The initial steady-state velocity for ATP hydrolysis (v_i_) was measured as a function of ATP concentration in buffer A at 4ºC [19]. The results for the PS/PS* complex and the P_2_S complex are shown in Supplemental **Figure S3-A** **and S3-B**, respectively. The data were fit to the Michaelis-Menten equation to obtain *k*_*cat*_ and K_m_ (**Table S3**, Supplemental **Figure S3**). These results show that both Rep(W250A) monomer (PS/PS*) and Rep(W250A) dimer (P_2_S) are able to hydrolyze ATP. The *k*_*cat*_ for wtRep and Rep(W250A) monomer are the same within error, although the K_m_ is ∼5-fold higher for Rep(W250A) monomer, consistent with the ∼10-fold lower affinity of Rep(W250A) for ssDNA. The *k*_*cat*_ for the Rep(W250A) dimer (P_2_S) is ∼5-fold lower than for wtRep dimer, whereas the K_m_ is only slightly higher (2.8 vs. 2.0 μM). In addition, Rep(W250A) also increased the K_m_ for PS/PS* complex by a factor of 5 while the that of P_2_S complex remained comparable. Thus, the W250A mutation has a dual effect on the ATPase activity. For PS complex, the mutation decreases the value of K_m_ but not the rate of ATP hydrolysis. Upon dimerization, for P_2_S, the mutation decreases the rate of hydrolysis but has no effect on the K_m_. Still, the Rep(W250A) protein is able to bind and hydrolyze ATP in a ssDNA-dependent manner.

### ssDNA translocation of Rep(W250A) and UvrD (W256A) monomers is impaired

Since Rep(W250A) monomers retain ATPase activity, we examined whether Rep(W250A) and UvrD(W256A) can undergo ATP-dependent directional translocation along ssDNA. We used a fluorescence stopped-flow method described previously for wtUvrD monomer [8, 9] and wtRep monomer [11] (**Figure 4**). This approach uses single strand DNA oligonucleotides of varying length, L, with a Cy3 fluorophore located either at the 5’ end, 5’-Cy3-(dT)_L_, or the 3’ end, (dT)_L_-Cy3-T-3’. Rep or UvrD is pre-mixed with an excess of DNA to ensure that they are bound as monomers. The enzyme will initially bind randomly along the ssDNA. For a translocase possessing 3’ to 5’ directionality, as is the case for Rep and UvrD, when the Cy3 is placed at the 5’ end of the DNA, any translocating protein that reaches the 5’ end will induce an enhancement in Cy3 fluorescence (PIFE) [56-59]. Upon addition of ATP, the resulting Cy3 fluorescence time course shows a transient increase in Cy3 fluorescence due to the translocases that reach the 5’ end, followed by a decrease due to protein dissociation from the 5’ end of the DNA. However, when the Cy3 is placed at the 3’ end, only a decrease in Cy3 fluorescence is observed due to translocation of any enzyme that happens to be bound to the 3’-end away from the 3’-end, as well as direct dissociation of enzyme from the ssDNA. The observed rate constant for this exponential decrease, k_obs_ = (k_t_+k_d_), and was used a constraint in global fits of the kinetic time courses (see **Scheme in Table 2**). A trap for free enzyme is added with the ATP to prevent rebinding of any free enzyme, thus ensuring a single round of translocation.

**Table 2.** Trp mutations weaken translocation.

|  | wtRep | Rep(W250A) | wtUvrD | UvrD(W256A) |
| --- | --- | --- | --- | --- |
| $k_t$ (steps/s) | $33 \pm 3$ | $1.5 \pm 0.1$ | $13.2 \pm 0.3$ | $0.5 \pm 0.2$ |
| $k_d$ (1/s) | 1.0(fixed) | 1.3(fixed) | $0.27 \pm 0.03$ | 1.0(fixed) |
| $k_{end}$ (1/s) | $0.08 \pm 0.001$ | $0.1 \pm 0.01$ | $0.02 \pm 0.001$ | $0.91 \pm 0.02$ |
| $k_c$ | $8.7 \pm 0.4$ | $18 \pm 7$ | $10.7 \pm 0.3$ | $63 \pm 341$ |
| $r$ | $0.20 \pm 0.03$ | $3.2 \pm 0.7$ | $0.52 \pm 0.02$ | $2.2 \pm 1.2$ |
| $c$ | $0.59 \pm 0.02$ | $0.5 \pm 0.2$ | $0.55 \pm 0.01$ | $3.4 \pm 217$ |
| $m$ (nt/step) | $7.2 \pm 1.3$ | $14.3 \pm 4.8$ | $8.0 \pm 0.1$ | $13.0 \pm 4$ |
| $P=k_t/(k_t+k_d)$ | 0.97 | 0.54 | 0.98 | 0.33 |
| $mk_t$ (nt/s) | $240 \pm 43$ | $22 \pm 7$ | $106 \pm 2$ | $7.0 \pm 2.0$ |

The final concentrations are 5 nM protein and 50 nM labeled DNA Cy3-(dT) . The buffer consists of 20 mM Tris pH 7.5, 6.0 mM NaCl, 10% v/v glycerol, 1 mM $\beta$ ME, 1.5 mM $MgCl_2$ , 0.5 mM ATP, 4 μM 3'-(dT)<sub>40</sub>-HP10 trap, at 25 °C.

**Figure 4.**
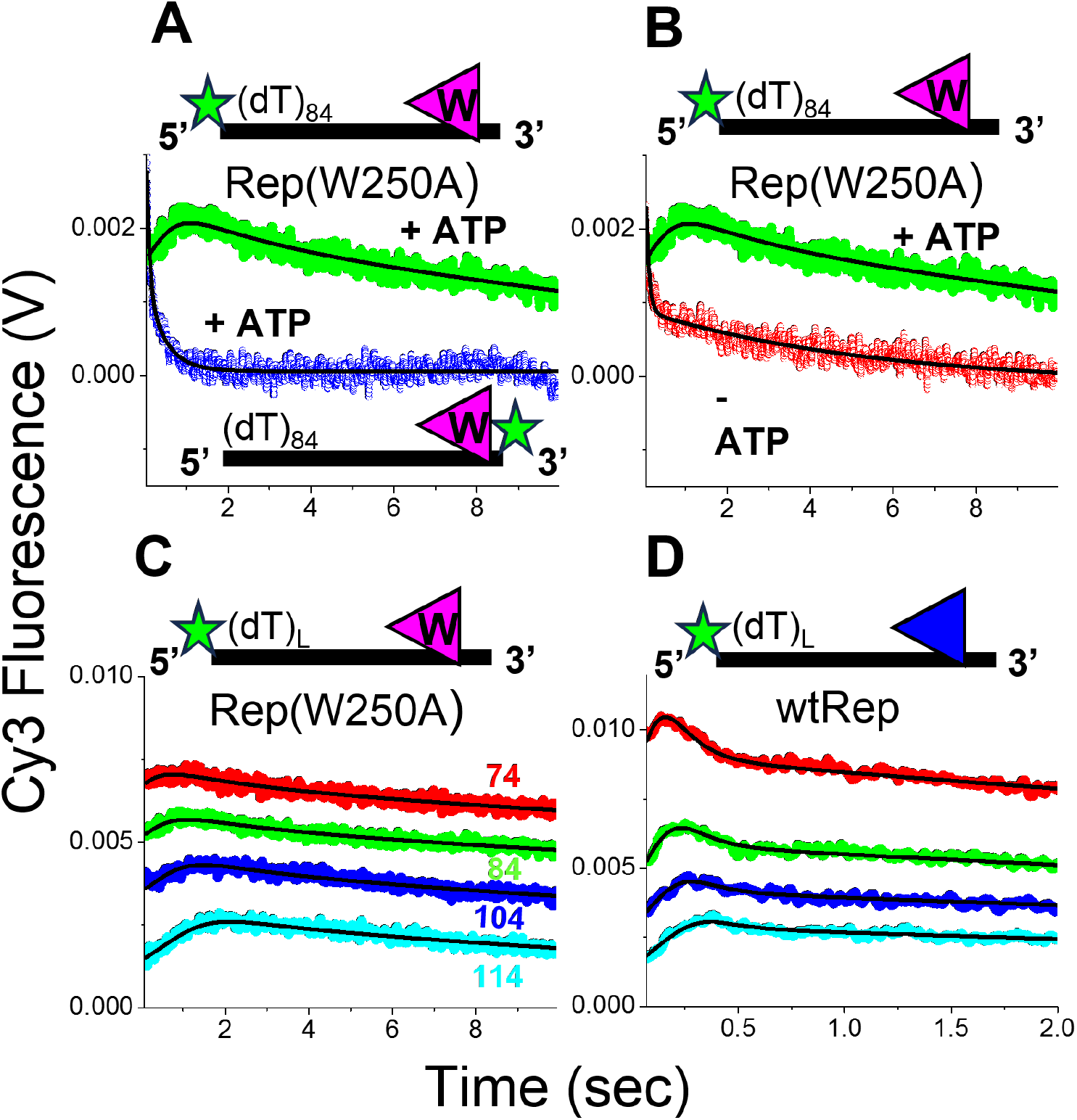
ATP-dependent translocation of Rep(W250A) on ssDNA. Stopped-flow fluorescence experiments were performed by mixing the contents of one syringe containing (100 nM DNA + 10 nM protein) with the contents of a second syringe containing (1.0 mM ATP, 3.0 mM MgCl_2_, 8 μM 3’-(dT)_40_-HP10 protein trap). (**A**)- (Green): Time course for Rep(W250A) on 5’-Cy3-(dT)_84_ showing that a Rep(W250A) monomer translocates toward the 5’ end of a ssDNA labeled with Cy3. (Blue): Time course for Rep(W250A) on ((dT)_84_-Cy3-T-3’) showing that a Rep(W250A) monomer translocates away from the 3’ end of a ssDNA labeled with Cy3. (**B**) No translocation is observed in the absence of ATP (red). (**C**) Time courses for Rep(W250A) monomer translocation on 5’-Cy3-(dT)_L_ with L = 74 (red), L = 84 (green), L = 104 (blue) and L = 114 (cyan). The traces are offset for clarity. The black lines are simulations based on global fits to **Scheme in Table 2** (see Methods) yielding a Rep(W250A) monomer translocation rate of 22 ± 7 nts/s (**Table 2**). (**C**) Time courses for wtRep monomer translocation on 5’-Cy3-(dT)_L_ with L = 74 (red), L = 84 (green), L = 104 (blue) and L = 114 (cyan). The traces are offset for clarity. Global fits yield a translocation rate of 240 ± 43 nts/s.

The results in **Figure 4A** indicate that Rep(W250A) monomers can translocate directionally along ssDNA. When the Cy3 is placed at the 5’-end of an oligodeoxythymidylate that is 84 nucleotides long (5’-Cy3-(dT)_84_), the time course upon addition of ATP shows the expected transient increase in Cy3 fluorescence indicating translocation in the 3’ to 5’ direction. However, when the Cy3 is placed at the 3’ end ((dT)_84_-Cy3-T), an exponential decrease in Cy3 fluorescence is observed upon addition of ATP indicating translocation away from the 3’-end. **Figure 4B** shows that translocation is ATP dependent, since only a decrease in Cy3 fluorescence is observed in the absence of ATP, reflecting Rep(W250A) dissociation from the DNA. Time courses for four different lengths of ssDNA, all with Cy3 at the 5’-end (5’-Cy3-(dT)_74,84,104,114_) (**Figure 4C**) show that the transient peak in the Cy3 fluorescence occurs at longer times as the length of the ssDNA is increased, as expected for an enzyme that translocates with 3’ to 5’ directionality. The time courses in **Figure 4C** were fit globally by non-linear least squares analysis to **Scheme in Table 2** to obtain the step rate constant for translocation, k_t_, and the rate constants for dissociation from internal sites, k_d_, the rate constant for dissociation from the 5’-end, k_end_, as well as the kinetic step size, m (see Methods). The macroscopic translocation rate is given by the product, mk_t_. The solid black lines in **Figure 4C** are simulations using Eq. (7) and the global fitting parameters given in **Table 2**. The macroscopic translocation rate for Rep(W250A) is 22 ± 7 nt/s (buffer U, 25 ºC). Under these same conditions, wtRep monomers translocate along ssDNA with a much faster macroscopic rate of 240 ± 43 nt/s (**Figure 4D**). In addition, the Rep(W250A) translocates with much lower processivity than wtRep (**Table 2**). Similar experiments performed with UvrD(W256A) (Supplementary **Figure S4**) show that it is also able to translocate directionally (3’ to 5”) along ssDNA, but also with a slower macroscopic rate of 7 ± 2 nt/s, compared to 106 ± 2 nt/s for wtUvrD under the same conditions (**Table 2**). Thus, although Rep(W250A) and UvrD(W256A) cannot unwind DNA, even as dimers, Rep(W250A) and UvrD(W256A) monomers are able to translocate along ssDNA, although translocation is quite impaired, being slower and less processive.

### Rep subunits communicate within a DNA bound Rep dimer

Moore and Lohman showed that ATP binding to Rep monomer, in the absence of DNA, does not affect Rep’s Trp fluorescence [60]. However, Hsieh *et al*. showed that ATP binding to a Rep dimer bound to ssDNA (dT_16_), elicits a complex change in Trp fluorescence as shown in **Figure 5A** [48]. Upon addition of ATP, Trp fluorescence undergoes an initial increase, followed by a decrease. The observed rates of the two Trp fluorescence changes both increase hyperbolically with ATP concentration, indicating that the two phases represent conformational changes induced by two ATP binding events, one to each Rep subunit. Therefore, these Trp fluorescence changes do not reflect ATP binding events, but rather conformational changes induced in the Rep dimer that follow ATP binding. In contrast, addition of the slowly hydrolyzable ATP-γ-S results in only the first Trp fluorescence increase, also hyperbolically dependent on ATP-γ-S [48]. Thus, the second Trp fluorescence change occurs only after hydrolysis of the first ATP bound to the Rep dimer. The P_2_S dimer formed with Rep(W250A) does not result in any Trp fluorescence change (**Figure 5A**), indicating that Trp250 is responsible for these fluorescence changes. Importantly, **Figure 5B** shows that no Trp fluorescence changes occur upon addition of ATP to wtRep monomer bound to ssDNA (dT)_16_. The Trp250 fluorescence changes induced by ATP reflect conformational changes that only occur within the ssDNA-bound Rep dimer, providing evidence for ATP-dependent communication between the two Rep subunits. We suggest that these ATP-dependent Rep dimer conformational changes are on the pathway to activating the Rep dimer helicase.

**Figure 5.**
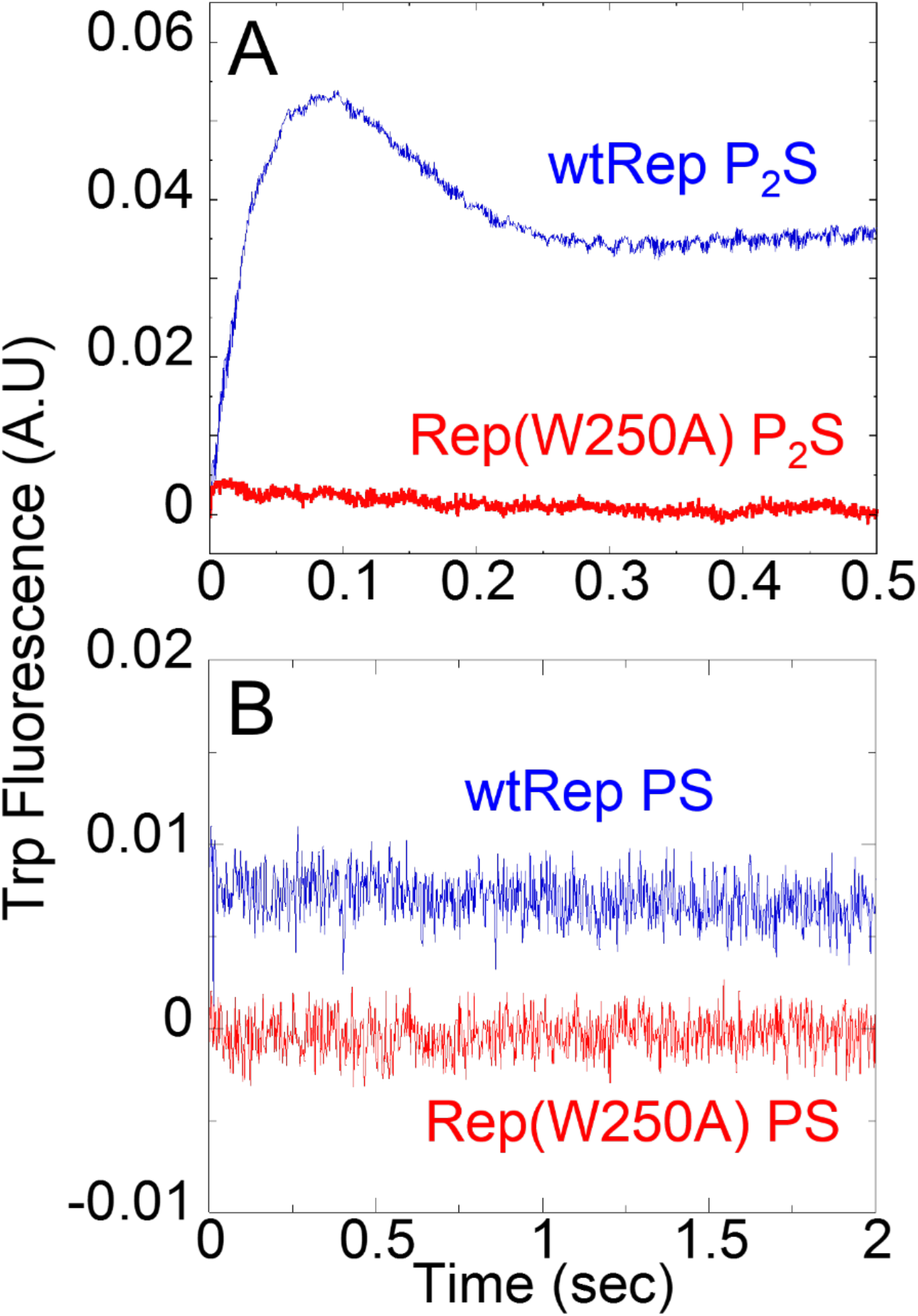
ATP binding causes a Trp fluorescence change only in a wtRep dimer bound to ssDNA (P_2_S), but not in a wtRep monomer bound to ssDNA (PS). Trp fluorescence was monitored using stopped-flow fluorescence upon mixing 200 μM ATP (buffer A, 4 ºC) with (**A**)- P_2_S dimer (800 nM wtRep or Rep(W250A) and 100 nM (dT)_16_), or (**B**) PS monomer (100 nM wtRep or Rep(W250A) and 5 μM (dT)_16_). Traces are offset for clarity.

### Rep(W250A) can activate the helicase activity of wtRep monomer

As shown previously, wtRep monomers possess ssDNA translocase activity, but not processive helicase activity. We also showed here that Rep(W250A) possesses ssDNA translocase activity, but not helicase activity, whether it is monomeric or dimeric. We next tested whether a hetero-dimer formed between Rep(W250A) and wtRep is able to activate helicase activity.

We first examined the ability of Rep(W250A) to activate a wtRep monomer helicase using ensemble rapid chemical quenched-flow experiments. wtRep (10, 20, 30, 50 nM) was premixed with an excess (60 nM) of DNA substrate (^32^P-labeled 3’-(dT)_20_-18-basepair duplex, see Methods) (Buffer U, 25 ºC), ratios where only wtRep monomers are bound. Each wtRep-DNA mixture was then mixed with Rep(W250A) and ATP. As shown in **Figure 6A**, in the absence of Rep(W250A), no DNA unwinding occurred at each wtRep concentration (10 - 50 nM) upon addition of ATP since only wtRep monomers are bound to the DNA under these conditions. However, DNA helicase activity is observed when the wtRep-DNA complex is mixed with Rep(W250A) and ATP. The extent of DNA unwinding increases with increasing Rep(W250A) titration. No unwinding occurs with Rep(W250A) in the absence of wtRep. The data suggest that Rep(W250A) can interact with wtRep and that the wtRep/Rep(W250A) hetero-dimer possesses helicase activity. The data for 10, 20 and 30 nM wtRep also show that the fraction DNA unwound first increases, but then decreases when mixed with a large excess of Rep(W250A) (>0.5 μM). This may be the result of Rep(W250A) displacing the wtRep or forming Rep(W250A) homo-dimers that are inactive.

**Figure 6.**
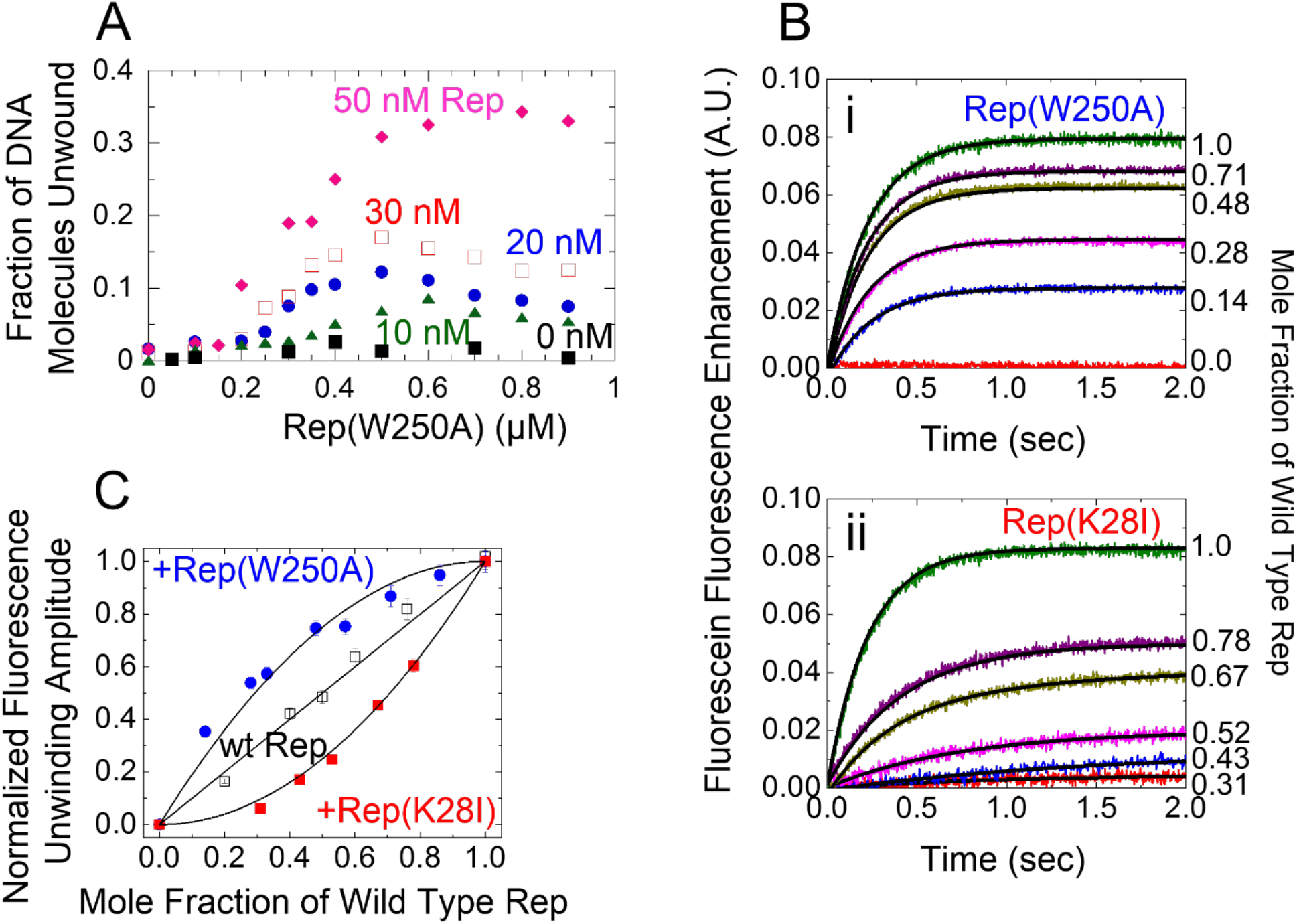
Helicase activity of wtRep monomer is activated by Rep(W250A) Rapid chemical quench flow experiments performed with an excess of DNA substrate (3’-(dT)_20_- 18bp: 60 nM) over wtRep at several concentrations (10nM, 20 nM, 30 nM and 50 nM) to ensure wtRep is bound as a monomer. No DNA unwinding is observed in the presence of wtRep alone (buffer U, 25ºC). The DNA unwinding reaction was initiated by addition of 1.5 mM ATP, 2 μM ssDNA (complementary to the unwound DNA product) to the pre- formed protein-DNA complex and a protein trap (5 μM 3’-(dT)_40_-HP10) to ensure single round conditions. DNA unwinding was stopped after 60 seconds with the addition of 0.4 M EDTA in 10% v/v glycerol. (**A**) Fraction of DNA unwound as a function of Rep(W250A) concentration. (**B**) Fluorescence stopped-flow traces showing DNA unwinding of fluorescein-hexachlorofluorescein labeled DNA (3’-(dT)_20_-18bp, F-HF) by (i) mixtures of wtRep and Rep(W250A) or (ii) mixtures of wtRep an ATPase deficient mutant, Rep(K28I) (buffer U, 25ºC). Experiments were conducted at 2 nM DNA + 100 nM total protein, varying the mole fraction of wtRep as indicated. The unwinding amplitude was determined by fitting the time course to a single exponential function. (**C**) DNA unwinding amplitudes, normalized to the unwinding amplitude for 100 nM wtRep, as a function of mole fraction of wtRep in the presence of Rep(W250A) (blue circles) or Rep(K28I) (red squares). Control experiments (black circles) were carried out with wtRep alone. For the Rep and Rep(W250A) experiment, the solid line is the prediction for three active dimeric species: wtRep:wtRep, wtRep:Rep(W250A), and Rep(W250A):wtRep (see Eq. (6)). For the wtRep and Rep(K28I) experiment, the solid line is the prediction for only one active species: wtRep:wtRep (see Eq. (5)).

Additional mixing experiments were conducted using a Forster resonance energy transfer (FRET) stopped-flow assay with a 3’-(dT)_20_-18-basepair DNA containing Fluorescein/Hexachlorofluorescein fluorophores at the blunt duplex end [61] (**Figure 6B**). DNA unwinding was monitored by the increase in Fluorescein fluorescence due to loss of FRET upon separation of the donor/acceptor pair. In these experiments, the DNA substrate concentration was 1 nM. The total protein concentration (wtRep + Rep(W250A)) was maintained constant at 100 nM, while the mole fraction of wtRep was varied. Hence, the protein is in large excess over the DNA. **Figures 6B-i** shows the DNA unwinding traces for the mixing with Rep(W250A) at different mole fractions of wtRep. Control experiments performed using an ATPase dead Rep mutant, Rep(K28I) [17] are shown in **Figure 6B-ii**. A second series of control experiments were performed in which only wtRep was included showing a linear dependence of the extent of DNA unwinding on wtRep concentration. The final amplitudes of the DNA unwinding traces are plotted as a function of wtRep mole fraction in **Figure 6C**. Addition of Rep(W250A) to wtRep (blue circles) showed an increase in the DNA unwinding amplitude relative to the wtRep alone control (black squares), while the addition Rep(K28I) to wtRep (red squares) showed a decrease in amplitude relative to wtRep alone.

These results (**Figure 6C**) indicate that wtRep/Rep(W250A) hetero-dimers possess helicase activity. The solid line describing the Rep(W250A) data is a simulation (Eq. (6)), based on the assumption that there are three active species in the population (wtRep/wtRep, wtRep/Rep(W250A), and Rep(W250A)/wtRep) and only the Rep(W250A) homo-dimer is inactive. These results also suggest that Rep(W250A)/wtRep heterodimers are active regardless of whether wtRep or Rep(W250A) is the lead subunit. The amplitudes would follow a straight line if only one of the hetero-dimers were active. Interestingly, the predicted curve for the Rep(W250A) data in **Figure 6C** is based on the assumption that the dimerization energetics of all three active dimers are equivalent (see Eq. (6) in Materials and Methods). As we have shown above, the Rep(W250A) dimer bound to ssDNA is ∼ 10-fold weaker than the wtRep dimer. However, the results of the mixing experiments suggest that the wtRep/Rep(W250A) hetero-dimers have energetic properties similar to wtRep homo-dimers. The solid line for the Rep(K28I) data is a simulation (Eq. (5)), based on the assumption that only the wtRep/wtRep homo-dimer has helicase activity, while both the Rep(K28I) homo-dimer and wtRep/Rep(K28I) hetero-dimers are inactive. These results indicate that a hetero-dimer formed between wtRep and Rep(W250A) can unwind DNA, whereas neither monomer alone has helicase activity. Furthermore, this activation by Rep(W250A) requires its ATPase activity.

Single Molecule experiments show helicase activity of Rep(W250A)/wtRep hetero-dimers and UvrD(W256A)/wtUvrD hetero-dimers.

The ensemble mixing experiments described above suggest that a Rep(W250A)/wtRep hetero-dimer possesses helicase activity. We considered whether the addition of Rep(W250A) to a wtRep monomer-DNA complex might result in some rearrangement of the wtRep on the DNA so that a wtRep homo-dimer was formed yielding helicase activity, although this could not lead to the shape of the mixing curve observed in **Figure 6C**. However, as an additional test we performed single molecule total internal reflectance fluorescence microscopy (smTIRFM) experiments designed to ensure that only one wtRep monomer remained on the DNA substrate upon mixing with Rep(W250A). For this experiment, N-terminal-biotinylated-wtRep monomer or N-terminal-biotinylated wtUvrD monomer were immobilized on a peg-neutravidin surface as described [28, 29](see Methods). A FRET labeled DNA substrate (3’-(dT)_20_-18 base pair duplex with a Cy3/Cy5 pair located on either end of the duplex region as in **Figure 7A**, see Methods section) was then added to bind to the surface bound wtRep or wtUvrD monomer. Rep(W250A) or wtRep was then added to the surface bound wtRep-DNA complex and ATP was added to initiate DNA unwinding (**Figure 7A**).

**Figure 7.**
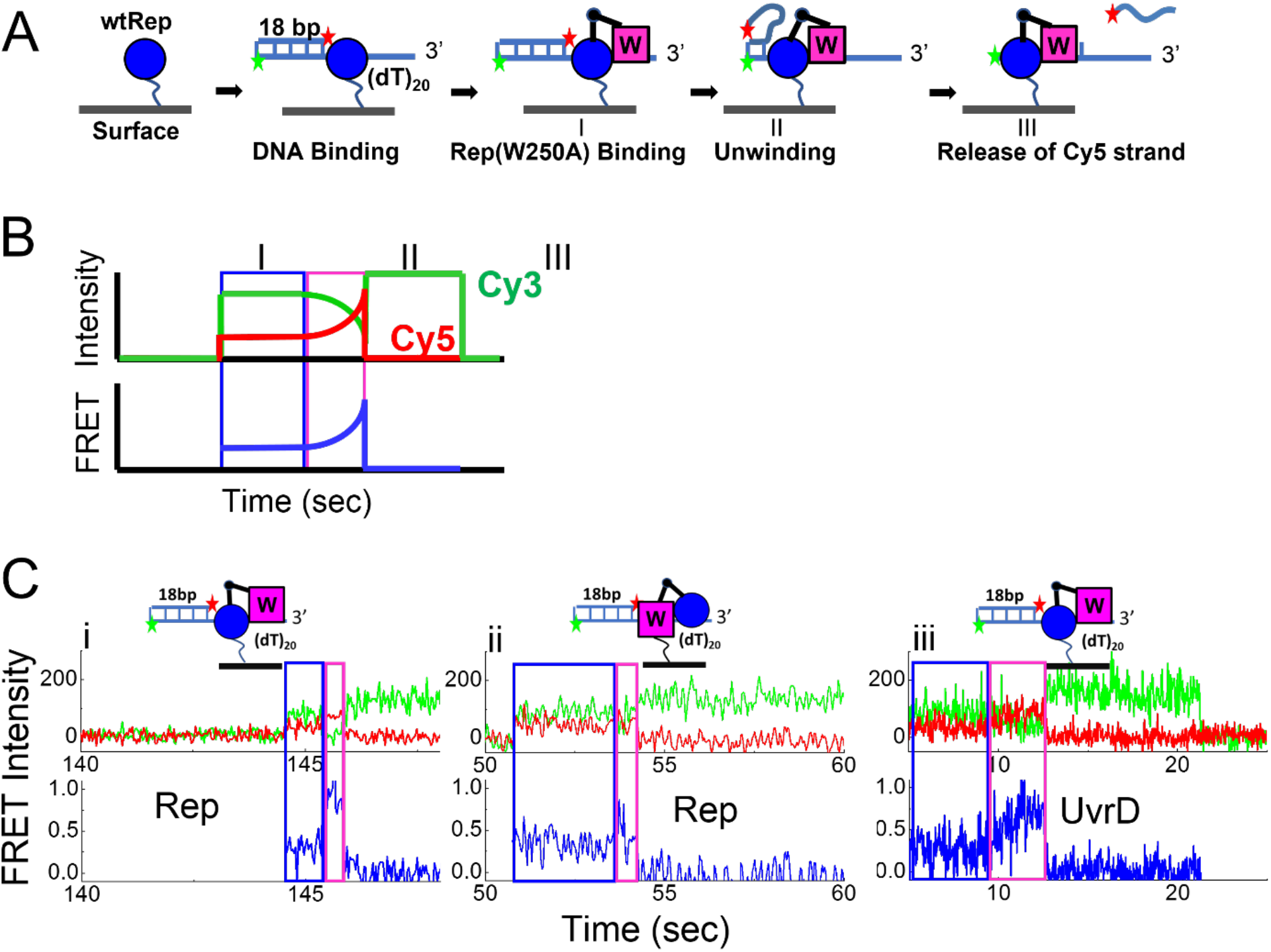
Single molecule TIRF experiments show activation of wtRep by Rep(W250A) and activation of wtUvrD by UvrD(W256A), but not cross activation. **(A)** Schematic of a single molecule FRET DNA unwinding experiment. N-terminal biotinylated monomeric proteins are immobilized on a PEG-NeutrAvidin surface. The reagents are added in the following order: 200 pM DNA substrate (3’-(dT)_20_-18bp) labeled with Cy3 and Cy5 as indicated, 400 nM Rep(W250A), and finally ATP (5-100 μM) to initiate the reaction. (**B**) DNA unwinding results in a transient increase in FRET followed by a loss of Cy5 fluorescence due to release of the top DNA strand. The DNA unwinding time (Δt (pink box)) is determined from the Cy3 and Cy5 traces. Unwinding rates were determined by dividing the duplex length (18bp) by the average unwinding time. (**C**) Examples of helicase activation of (i) biotin-wtRep + Rep(W250A) at 50 μM ATP; (ii) biotin-Rep(W250A) + wtRep at 50 μM ATP, and (iii) biotin-wtUvrD + UvrD(W256A) at 25 μM ATP. Experiments were performed in SM buffer at 25 ºC.

In this assay, DNA unwinding is observed as an anti-correlated change in Cy3 and Cy5 fluorescence resulting in a transient increase in FRET followed by a loss of Cy5 fluorescence and an increase in Cy3 fluorescence as depicted schematically in **Figure 7B**. The DNA unwinding time, Δt, is defined as the time from where the FRET signal starts to increase until the Cy5 signal disappears (**Figure 7B**). Examples of Rep or UvrD heterodimer unwinding are shown in **Figure 7C** for biotin-Rep:Rep(W250A), biotin-Rep(W250A):Rep, and biotin-UvrD:UvrD(W256A). The DNA unwinding rates were calculated from the average DNA unwinding time (n/k) and the unwinding length (18 base pairs) (Eq. (8) [28, 29, 39, 62, 63]. The [ATP] dependence of the DNA unwinding rates was fit to the Michaelis-Menten equation yielding V_max_ and K_m_ (**Figures 8A** (Rep) and **8B** (UvrD), **Table 3**). As shown in **Table 3**, both wtRep/Rep(W250A) hetero-dimers and wtUvrD/UvrD(W256A) hetero-dimers unwind ∼two-fold slower than the respective homo-dimers. We also performed experiments in which biotinylated-Rep(W250A) is immobilized on the surface, followed by addition of labeled-DNA and wtRep (**Figure 7C-ii**), but found the same rates of unwinding for the hetero-dimers regardless of which monomer was surface bound (**Table 3**). The percentage of molecules unwound is the same for homodimer as for heterodimer Rep (17-27%) across three ATP concentrations (**Table S4**). For UvrD, the percentage of molecules unwound for the heterodimers is lower than of the homodimers at high ATP concentrations (**Table S5**). In the absence of added proteins, no DNA unwinding was observed by the surface bound protein indicating that the immobilized monomers cannot unwind DNA. Importantly, no cross activation between Rep and UvrD was observed (<1%) although wtUvrD and wtRep can form a heterodimer [64]. That is, wtRep monomer was not activated by wtUvrD or UvrD(W256A) and wtUvrD was not activated by wtRep or Rep(W250A) (**Table 3**). Hence activation involves specific interactions with both the Rep dimer and UvrD dimer.

**Table 3.** Helicase activation requires specific dimer interactions.

| **Added** | **wtRep**<br>$V_{\max}$ (bp/s)<br>$K_M$ (μM) | **Rep(W250A)**<br>$V_{\max}$ (bp/s)<br>$K_M$ (μM) | **wtUvrD**<br>$V_{\max}$ (bp/s)<br>$K_M$ (μM) | **UvrD(W256A)**<br>$V_{\max}$ (bp/s)<br>$K_M$ (μM) |
| --- | --- | --- | --- | --- |
| **Immobilized** |  |  |  |  |
| **biotin-Rep** | 49.3 ± 0.1<br>12.8 ± 0.1 | 29.7 ± 0.4<br>8.4 ± 0.4 | No Activation | No Activation |
| **biotin-RepW250A** | 26.0 ± 0.4<br>5.6 ± 0.3 | No Activation | No Activation | No Activation |
| **biotin-UvrD** | No Activation | No Activation | 43.8 ± 2.3<br>12.6 ± 1.5 | 28.9 ± 1.5<br>8.0 ± 1.1 |

**Figure 8.**
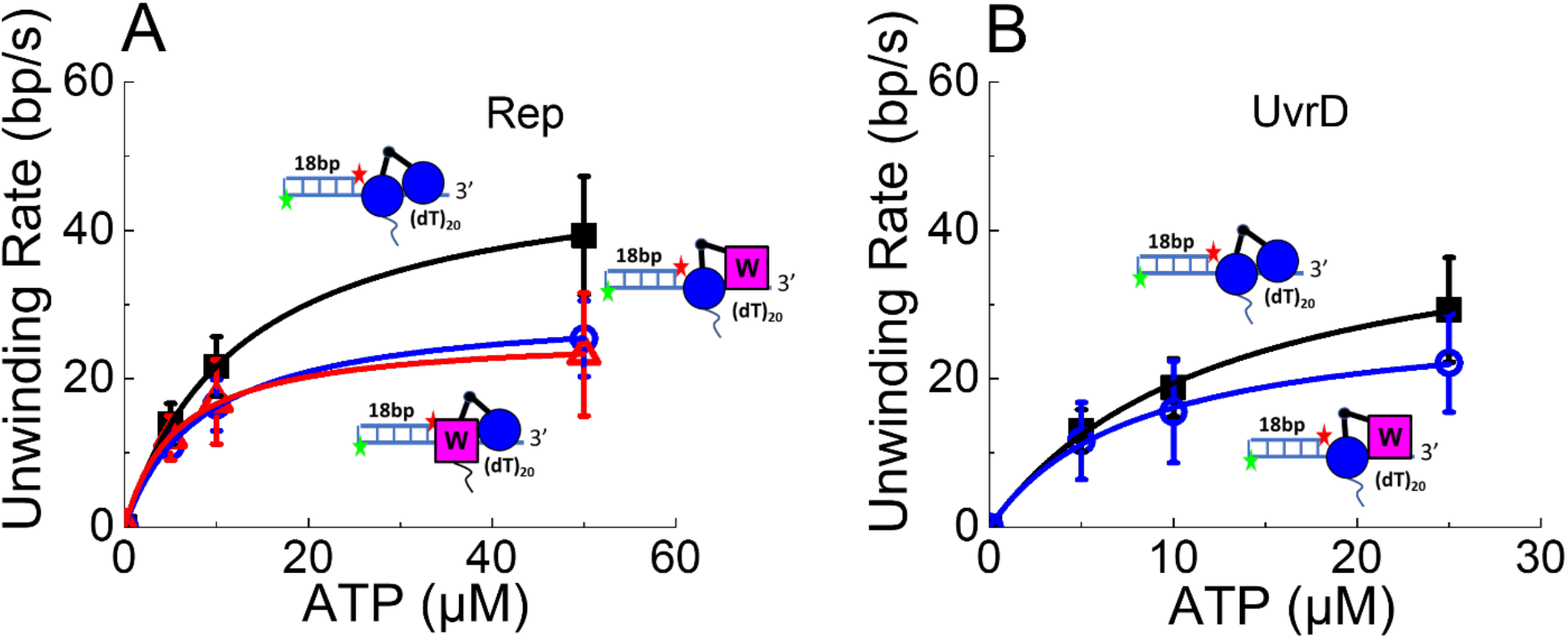
ATP dependence of DNA unwinding rates from smTIRF experiments. (**A**) wtRep homo-dimers unwinding rates are higher than for wtRep:Rep(W250A) heterodimers regardless of which subunit is immobilized on the surface. (**B**) wtUvrD homo-dimers unwind with higher rates than wtUvrD:UvrD(W256A) hetero-dimers. Solid lines are best fits to the Michaelis-Menten equation (see **Table 3**). The data at every ATP concentration is the average unwinding rate (18 bp / average unwinding time (Δt)) for 14-34 individual molecules (Tables S4 and S5). Error bars represent one standard deviation. Experiments were conducted in SM buffer at 25 ºC.

## Discussion

We have shown for both Rep and UvrD that a gain of helicase activity can be achieved by hetero-dimerization of one wild type monomer and one Trp-mutant monomer (Rep(W250A) or UvrD(W256A)), neither of which possess helicase activity as monomers. Importantly, the Trp-mutant proteins can still dimerize, but do not possess helicase activity even when dimeric. The Rep(W250A) and UvrD(W256A) monomers also retain ATP-dependent ssDNA translocase activity, but this activity is greatly impaired compared to the wt monomers. Specific interactions are required within each hetero-dimer to induce helicase activity since neither wtRep nor Rep(W250A) can activate a wtUvrD monomer and neither wtUvrD nor UvrD(W256A) can activate a wtRep monomer. Furthermore, an ATPase dead Rep mutant, Rep(K28I) is unable to activate wtRep monomer, indicating that the second activating subunit needs to hydrolyze ATP, which is the case for the Rep(W250A) and UvrD(W256A). The rate of DNA unwinding by the hetero-dimers is ∼two-fold slower than for the wt homo-dimers. Since the Rep(W250A) and UvrD(W256A) monomers have much slower rates of ssDNA translocation compared to wtRep or wtUvrD, this suggests DNA unwinding by the hetero-dimers may require the translocase activity of the Trp-mutant subunit.

There is abundant evidence that regulation of the helicase activity of Rep, UvrD, and PcrA involves the auto-inhibitory 2B sub-domains [11] that can undergo substantial rotation with respect to the remaining sub-domains in all three enzymes [28, 41-45, 65]. Activation of the UvrD helicase correlates with movement of the 2B sub-domain both by dimerization [28] or by interaction with the accessory protein, MutL [39, 40]. However, cross reactivity between Rep and UvrD does not occur as Rep is unable to induce movement of the UvrD 2B domain and does not activate helicase activity of the UvrD monomer [28] and vice versa, as we show here. Furthermore, when a DNA substrate is under tension, UvrD monomer can display limited helicase activity that correlates with a “closed” conformation of the 2B sub-domain [65]. Rep monomer helicase activity can also be activated by deleting the auto-inhibitory 2B sub-domain [11, 31, 32] or by crosslinking of the 2B sub-domain into a closed conformation [33].

We have hypothesized that homo-dimerization of Rep and also of UvrD and PcrA involves direct interactions between the 2B sub-domains of each subunit [41] and that dimerization results in movement of the 2B sub-domains that relieves the auto-inhibition [1, 11]. This hypothesis is supported by the recent finding that the homologous SF1A helicase, UvrD1, from *Mycobacterium tuberculosis* (*Mt*) also requires dimerization for helicase activity. *Mt* UvrD1 dimerization occurs via a redox-dependent Cystine crosslink between the 2B sub-domains of each subunit [46]. Such a 2B-2B interaction also occurs between the RecB and RecC subunits of the RecBCD helicase [66]. Crystal structures of monomers of PcrA [43] and UvrD [44] bound to a short duplex DNA possessing a 7 nucleotide flanking 3’-ssDNA show the 2B sub-domain interacting with the DNA duplex. Even though those DNA substrates cannot be unwound by monomers, or even dimers of PcrA or UvrD, it has been proposed that monomers are the active forms of these helicases and that 2B sub-domain duplex interactions are essential for helicase activity [43, 44]. Yet, Rep monomer helicase is activated by deletion of its 2B sub-domain [11].

The conserved aromatic amino acid (Trp for Rep and UvrD) is required in at least one subunit of the dimer for helicase activity. Dimerization must alter the DNA binding site which contains the Trp residue. By linkage, the DNA binding site must also alter dimerization energetics. This is certainly the case for Rep since Rep dimerization requires DNA binding [12-14, 51, 60, 67]. In addition, Rep monomers and Rep dimers have vastly different intrinsic ssDNA-dependent ATPase activities [17-19, 48] indicating that dimerization and ATPase activity are energetically linked. Furthermore, we show here that the Rep W250A mutation affects Rep dimer formation and dimer stability.

Additional evidence that Rep dimerization and ssDNA binding are linked comes from the observation that ATP binding and hydrolysis results in a Trp250 fluorescence change only in the ssDNA bound Rep dimer, P_2_S [48], but not free Rep monomer [60, 67] or Rep monomer bound to ssDNA (PS), as we show here. Hence, Rep dimerization must induce a conformational change within the DNA binding sites. If dimerization of Rep occurs via the 2B sub-domains, as is the case for *Mt* UvrD1, this suggests long range allosteric communication between the dimerization domains and the DNA binding and ATP binding sites.

Crystal structures indicate that the conserved Trp residue in motif III (Trp-250 in Rep and Trp-256 in UvrD and Trp-259 in PcrA) forms a stacking interaction with the DNA bases [41, 43, 44]. Removal of the stacking interaction by substituting Trp for Ala likely contributes to the loss of helicase activity. However, the resulting mutants, Rep(W250A) and UvrD(W256A) still bind DNA, dimerize and hydrolyzes ATP, and monomers can still translocate along ssDNA, although each of these activities is compromised severely. The importance of this conserved aromatic amino acid in Motif III within the DNA binding site has been demonstrated for other helicases. *Mycobacterium* AdnAB is a hetero-dimeric SF1 helicase involved in double strand break repair. The AdnB subunit has ATPase activity and contains a Trp residue (W325) in motif III within its ssDNA binding site. Warren *et al*. [68] made several mutations at position 325 and showed that DNA unwinding activity was maintained as long as an aromatic amino acid replaced W325. However, mutation to Ala or Leu abolished DNA unwinding activity.

### Subunit switching models

Two subunit switching models have been proposed for how a dimeric helicase might function: a rolling model [69] and an inch-worm model [70]. The major difference between the two models is that in the inch-worm model the same subunit remains in the lead position (**Figure 9**), whereas in the rolling model, the two subunits alternate as the lead subunit [69], similar to the hand-over-hand models for dimeric kinesin translocation along microtubules [71, 72]. Although we currently favor the dimeric inch-worm model, we are unable to rule out the rolling model at this point. The fact that a monomer that is severely impaired for DNA binding, ATPase and ssDNA translocase activity is able to dimerize with a wt subunit and activate the helicase suggests that both subunits do not need to possess helicase activity. If the helicase activity resides in the wt subunit then this suggests that the wt subunit would be maintained as the lead subunit in the asymmetric Rep(W250A)/wtRep and UvrD(W256A)/wtUvrD hetero-dimers, which would support a dimeric inch-worm model. However, since the properties and activities of the individual subunits are clearly modified by dimerization, it is possible that either subunit can serve as the lead helicase. In fact, our single molecule TIRFM experiments show the same DNA unwinding rates and the percentage of molecules unwound regardless of whether the wild type monomer or the Trp-mutant monomer is first immobilized on the surface, suggesting that either subunit could serve as the lead subunit. This conclusion is also consistent with the stopped-flow mixing experiments which suggest that Rep(W250A)/wtRep hetero-dimers have helicase activity regardless of which is the lead subunit.

**Figure 9:**
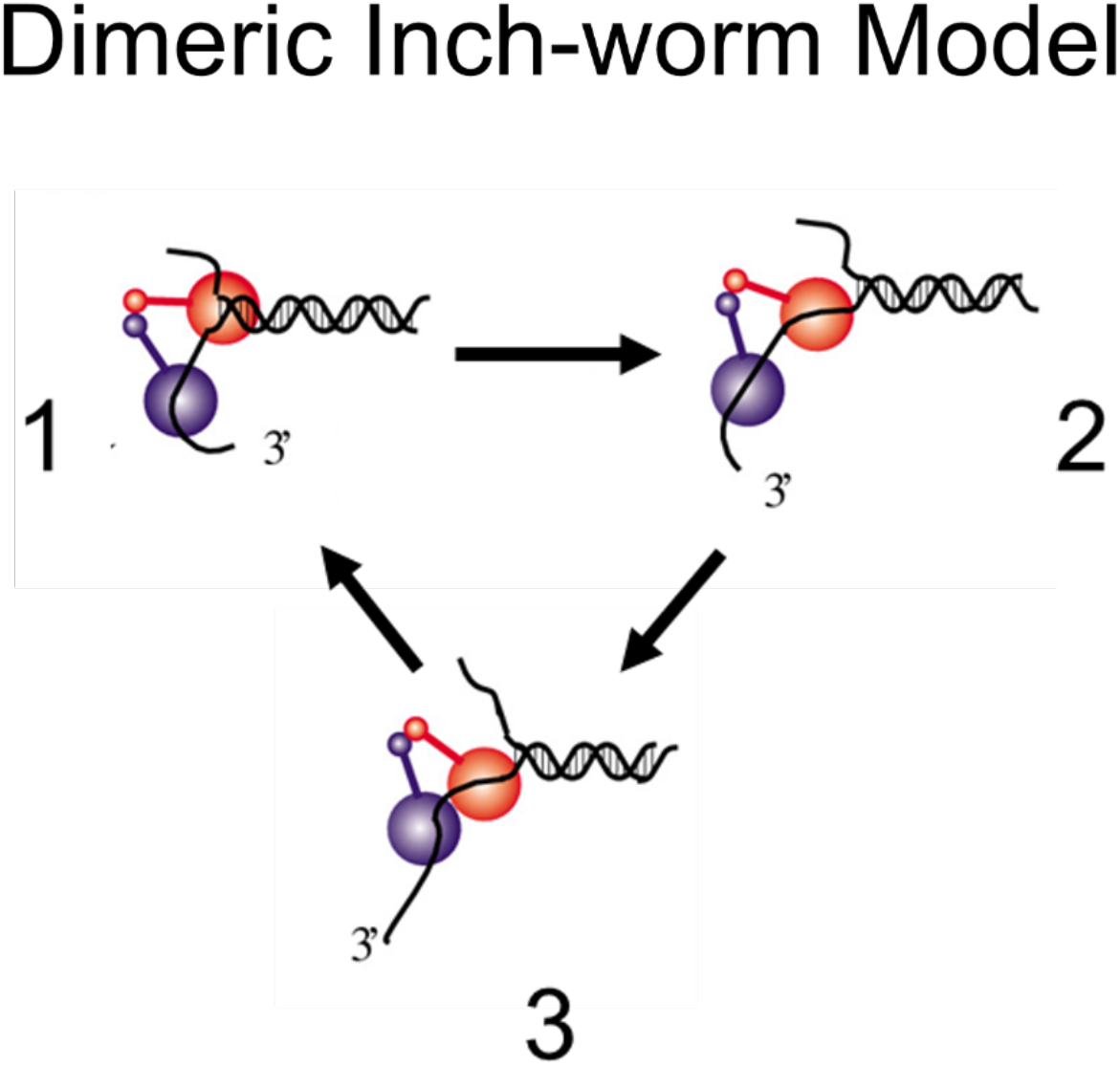
Dimeric Inch-worm model for DNA unwinding. One possible model for DNA unwinding by a dimeric helicase in which the lead subunit remains the lead subunit. State (1)- the lead subunit (red) binds to the ss/ds DNA junction. State (2)- the lead subunit melts some number of base pairs within the duplex DNA at the ss/ds DNA junction. State (3)- the trailing subunit (blue) translocates along the ssDNA to reset the helicase. In a dimeric rolling model [64] the lead and trailing subunits would switch after each cycle.

It is important to note that the helicase activities of the SF1 enzymes, *E. coli* Rep, *E. coli* UvrD, and *B. stearothermophilus* PcrA can also be activated by other mechanisms in addition to dimerization. A Rep monomer can be activated by interaction with PriC [29], and a UvrD monomer can be activated by MutL [39, 40], although neither of these accessory proteins have translocase activity on their own. Those accessory proteins presumably relieve the auto-inhibitory effects of the 2B sub-domain through their interactions as well, but the mechanism of DNA unwinding may differ in those cases.

As a final point, we emphasize that the molecular basis for helicase “activation” and the mechanisms of DNA unwinding are not understood for any SF1 helicase. Mechanistic evidence [1] indicates that helicase activity does not occur via the mechanical models that have been postulated based on the monomeric structures of UvrD and PcrA [43, 44]. This is underscored by the fact that the monomeric forms of UvrD, PcrA and Rep are not active helicases on their own in the absence of force applied to the DNA. One possibility is that activation enables the helicase to actively destabilize (“melt”) some number of base pairs simply upon binding to the ss/dsDNA junction in order to initiate DNA unwinding as is the case for RecBCD [66, 73-77]. Both the mechanisms of activation and DNA unwinding by SF1 helicases remain outstanding questions.

## Materials and Methods

### Buffer and reagents

Buffer A is 20 mM Tris pH 7.5, 6.0 mM NaCl, 5.0 mM MgCl_2_, 10% v/v glycerol. Buffer U is 20 mM Tris pH 7.5, 6.0 mM NaCl, 1.5 mM MgCl_2_, 10% v/v glycerol. The SM buffer (for single molecule experiments) is buffer U + 3.0 mM Trolox, 20 units/ml catalase, 20 units/ml glucose oxidase, 0.8% w/v dextrose, and indicated ATP concentrations.

### Proteins and DN

Rep and UvrD were purified as described [15, 78]. Rep and UvrD proteins containing a biotin tag at the N-terminus were prepared as described [28, 29]. Oligodeoxynucleotides were synthesized on a MerMaid 4 DNA synthesizer (BioAutomation Corporation, Irving, TX) and further purified by gel-electrophoresis and HPLC (Waters Corporation, Milford, MA) as described [79].

### ΦX174 replication assay

Bacteriophage φX174 replication in *E. coli CK11Δrep/pIWcI* in different *rep* genetic backgrounds was examined using a standard plaque assay at 37 ºC [53]. ΦX174 was purchased from ATCC (Manassas, VA). Control experiments with *CK11Δrep/pIWcI* ± pRepO were included in each plaque assay. Steady-state ATPase activity of wtRep and Rep(W250A) were determined in Buffer A at 4 ºC as described [17, 19].

### Single time point determination of Rep unwinding amplitude

The ^32^P-labeled 3’- (dT)_20_-ds18 DNA (60 nM) was incubated with wtRep (50 nM) to initially form a Rep monomer-DNA complex, followed by the addition of wtRep or mutant Rep (0 to 1 μM), in Buffer U (50 μl) at 25 ºC for 5 minutes. DNA unwinding was initiated by adding an equal volume of 2 mM ATP and 5 μM protein trap. The unwinding reaction was stopped after 60 seconds by adding 200 μl of 0.4 M EDTA + 10% v/v glycerol. The unwinding products were analyzed by gel electrophoresis and the fraction of unwound DNA was determined as described above.

### Equilibrium fluorescence titration of ssDNA with Rep

Equilibrium fluorescence titrations of 2-aminopurine (2-AP)-containing ssDNA (5’-(dT)_5_-(2-AP)-(dT)_4_-(2-AP)-(dT)_5_) were performed in Buffer A as described [51]. The 2-AP was excited at 315 nm (2 mm slits), and fluorescence was monitored at λ_em_ ≥ 350 nm (using a long pass Oriel filter).

### Ensemble DNA unwinding kinetics

Single round rapid chemical quenched-flow DNA unwinding experiments were conducted as described [80] using a KinTek RQF-3 rapid chemical quenched-flow apparatus (KinTek Crop, Austin, TX). The DNA unwinding substrate consisted of an 18 base pair duplex with a 3’-(dT)_20_ tail. The top strand was labeled with ^32^P at the 5’ end by T4 polynucleotide kinase (USB Corp, Cleveland, OH). A DNA hairpin 3’-(dT)_40_-HP10 (5 μM) was used as a protein trap to ensure a single round of DNA unwinding.

DNA unwinding experiments were performed using a DNA unwinding substrate (1 nM final concentration) at different mole fractions of wtRep, at a constant total protein concentration of 100 nM (final concentration).

Single round fluorescence stopped-flow experiments were conducted with either a SX-17MV (with a 150W Xe Arc lamp) or a SX-20 stopped-flow instrument (with an LED source) (Applied Photophysics, Leatherhead, Surrey, UK) as described [39, 40]. The 3’- (dT)_20_ with 18-bp double strand DNA was labeled at the blunt ends with a pair of Cy5 and a black hole quencher (BHQ2) or fluorescein, F and hexachlorofluorescein, HF (donor and acceptor). The labeled-DNA/protein was pre-incubated in one syringe and mixed with ATP + protein trap in the second syringe [61]. In the wtRep: mutant Rep mixing experiments, wtRep and mutant Rep (Rep(W250A) or Rep(K28I)) were mixed in different wtRep mole fractions (final protein concentration maintained at 100 nM) and preincubated with fluorescent unwinding substrate (2 nM of 3’-(dT)_20_-18bp dsDNA labeled with H/HF pair; λ_ex_ = 492 nm, λ_em_ ≥ 520 nm) in buffer U 25 ºC for 5 minutes. The unwinding reaction was initiated by rapid mixing of the complex with 2 mM ATP and competitor DNA ((dT)_16_, 5 μM, ε_260_ = 1.30 × 10^5^ M^−1^ cm^−1^). The presence of the protein trap, (dT)_16_, prevented the free Rep from rebinding to the unwinding substrate and reinitiating the unwinding, thus, maintaining the single-turnover conditions. The fluorescence time courses were fit to either a single or the sum of exponential terms, n, using Eq. (1).

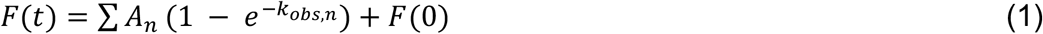

Where F(t) is the total fluorescence amplitude at time t, n is the number of exponential terms, and A_n_ and k_obs,n_ are the amplitude and the observed rate constant for each term, respectively. *F*(0) is the fluorescence at time zero. The fluorescence traces shown in the figures represent an average of 6-14 individual determinations, and the indicated errors in the fitted parameters are the standard errors.

### Stopped-flow Rep hetero-dimer mixing experiments to examine wtRep/Rep(W250A) activity

In a mixing experiment containing wtRep and mutant Rep (either Rep(W250A) or Rep(K28I)), we define the mole fraction of wtRep as *a* = [wt]/([wt] + [mutant]) and the mole fraction of mutant Rep as *b =* [mutant]/([wt] + [mutant]). The probability of finding a wtRep in the mixture is ρ_W_ = *a*/(*a +b)*, and the probability of finding a mutant Rep in the mixture is ρ_M_ = *b*/(*a +b)*. If the dimerization energetics of the wtRep and Rep mutant are equivalent, then the probabilities of forming the four different Rep dimers will be given by [18]:

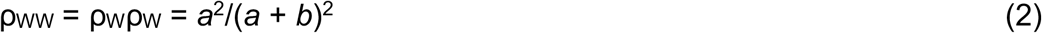

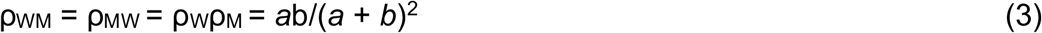

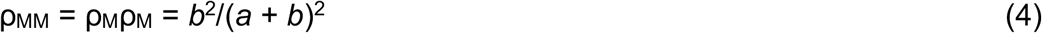

The normalized total amplitudes, A_T_ from the single round stopped-flow fluorescence DNA unwinding experiments containing a mixture of wtRep and Rep mutant (either Rep(W250A) or Rep(K28I)) were determined as a function of the mole fraction of wtRep protein. The wtRep homo-dimer has full DNA helicase activity and the Rep(W250A) homo-dimer has no helicase activity. Therefore, as shown previously [18], if the wt/mutant hetero-dimer inhibits DNA unwinding activity, then only the wild-type Rep homodimers will be active, and the total DNA unwinding amplitude in a single round DNA unwinding experiment, A_T_, will be described by Eq. (5), where *a* = [wt]/([wt] + [mutant]) is the mole fraction of wtRep and *b* = [mutant]/([wt] + [mutant]) is the mole fraction of mutant Rep. If the hetero-dimeric wt/mutant protein complex possesses DNA unwinding activity, then three dimers will be active (Rep:Rep, Rep:mutant Rep, mutant Rep:Rep) and the total unwinding amplitude in a single round described by Eq. (6).

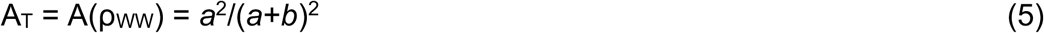

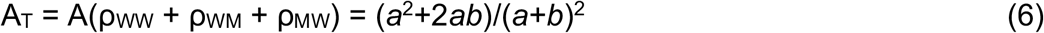

For unwinding experiments using Cy5/BHQ2 DNA, the fluorescence was monitored by exciting Cy5 with a 625-nm LED (18 mA), and the fluorescence emission was passed through an emission filter 665-nm long pass filter (Oriel, Stratford, CT) using a SX-20 fluorescence stopped-flow instrument. The protein and DNA were premixed in Buffer U at 25 ºC (having 1.5 mM MgCl_2_) in one syringe while the other syringe contains ATP and a protein trap in the same buffer. The trap for free protein (3’-(dT)_40_-HP10) was included to prevent the rebinding of the protein to DNA and thus ensure single-round kinetics. The increase in Cy5 fluorescence indicates the DNA unwinding as the Cy5 is separated from the quencher BHQ2. [28, 29]

The stopped-flow ssDNA translocation experiments were conducted with Cy3 labeled DNA on the 5’ end, Cy3-(dT)_L_. DNA-protein solution (100 nM DNA + 10 nM protein) was initiated by mixing with 3.0 mM MgCl_2_, 1.0 mM ATP and 8.0 μM of a protein trap (3’-(dT)_40_-HP10) in buffer U at 25 ºC. The large excess of DNA over the protein to ensure that a monomer is bound to the DNA. The sample was excited with a 505-nm LED, and the fluorescence emission was filtered with a 520-nm long pass filter. The fluorescence profiles were fitted to a n-step translocation model (**Scheme in Table 2**, Eq. (7)) to obtain the macroscopic translocation rates as previously described. [7-10, 81]

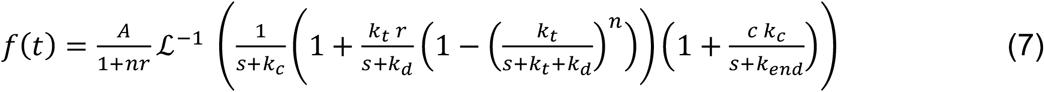

*A* is the total protein concentration bound at time zero; r is the ratio of protein concentration bound and any position other than 5’-end to the protein at the 5’-end at time zero; and *c* (= f^*^_end_/f_end_) is the ratio of fluorescence signals in states *I*_*end*_ and *I*^*\**^_*end*_.

### Single molecule FRET

The SM experiments were conducted with an Olympus IX71 TIRF microscope equipped with a 60x Olympus TIRF objective as described [28, 29]. All experiments were conducted at 25 ºC. The DNA labeled with Cy3 fluorophore was excited with a 532-nm laser, and the fluorescence emissions were passed through a dual image splitter and recorded with an Andor iXon Ultra 897 EMCCD camera (Oxford Instruments) [79]. The data were collected at 31 Hz and processed with IDL and MATHLAB scripts [82]. For DNA unwinding experiments, the biotinylated helicases (Rep or UvrD, 50 pM) were immobilized on the neutravidin surface for 5 minutes and then washed to remove unbound protein and then Cy3/Cy5 labeled DNA substrate (10 μl of 0.2 nM) was added and incubated for 1 min. Untagged-proteins (400 nM) were then added and incubated for 3 minutes. DNA unwinding was initiated by adding ATP. The time-dependent changes in Cy3 and Cy5 fluorescence, and subsequently the FRET trace were analyzed to obtain the unwinding time (Δt), the period from the FRET increasing to the dissociation of Cy5, and the average unwinding time (n/k) (Eq. (8) was used to calculate the average DNA unwinding rate as described [28, 29, 39, 62, 63],

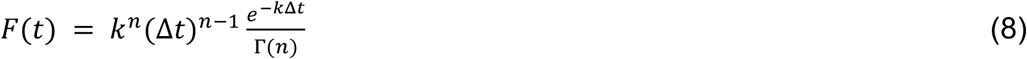

where *k* is the stepping rate (steps/s) and *n* is the number of steps. Since *k* = *k*_*u*_ + *k*_*d*_ (where *k*_*u*_ = unwinding rate and *k*_*d*_ = dissociation rate), *k* ∼ *k*_*u*_ when *k*_*u*_ >> *k*_*d*_. The DNA unwinding rate, r = L/(n/k), where L is the unwinding length in bp and (n/k) is the average unwinding time.

## Acknowledgements

We thank Kacey Mersch, Roberto Galletto, Eric Tomko, Ankita Chadda and Eric Galburt for helpful discussions, Elizabeth Weiland, Thang Ho and Bill van Zante for technical assistance. This research was supported in part by the NIH (R35 GM136632 and RO1 GM45948 to TML).

## Supplementary materials

### Materials and Methods

The concentrations of DNA were determined using their extinction coefficients per strand. The extinction coefficients for other molecules are: Fluorescein (ε_495_ = 7.5 × 10^4^ M^−1^ cm^−1^), Hexachlorofluorescein (ε_538_ = 9.6 × 10^4^ M^−1^ cm^−1^), Cy3 (ε_260_ = 5.0 × 10^3^ M^−1^ cm^−1^, ε_550_ = 1.37 × 10^5^ M^−1^ cm^−1^), Cy5 (ε_260_ = 1.0 × 10^4^ M^−1^ cm^−1^, ε_650_ = 2.5 × 10^5^ M^−1^ cm^−1^), BHQ2 (ε_260_ = 8.0 × 10^3^ M^−1^ cm^−1^, ε_579_ = 3.8 × 10^4^ M^−1^ cm^-1^), ATP (ε_259_ = 1.54 × 10^4^ M^−1^ cm^−1^), Trolox (ε_290_ = 2350 M^−1^ cm^−1^). MgCl_2_ stock concentrations in water were determined by <u>refractive index</u> measurements at 20 °C (ABBE Mark II Refractometer) using the reference data from the CRC handbook.

The protein concentrations were determined spectrophotometrically in buffer A using their extinction coefficients (Rep, ε_280_ = 7.68 × 10^4^ M^−1^ cm^−1^; Rep(W250A), ε_280_ = 7.13 × 10^4^ M^−1^ cm^−1^; Rep(K28I), ε_280_ = 7.68 × 10^4^ M^−1^ cm^−1^; UvrD, ε_280_ = 1.06 × 10^5^ M^−1^ cm^−1^; UvrD(W256A), ε_280_ = 1.00 × 10^5^ M^−1^ cm^−1^).

For the fluorescence stopped-flow ssDNA translocation experiments, the extinction coefficients for the DNA oligomers Cy3-(dT)_L_ are: L=74: ε_260,dna_ = 6.00 × 10^5^ M^−1^ cm^−1^, L=84: ε_260,dna_ = 6.81 × 10^5^ M^−1^ cm^−1^, L=104: ε_260,dna_ = 8.43 × 10^5^ M^−1^ cm^−1^, and L=114: ε_260,dna_ = 9.24 × 10^5^ M^−1^ cm^−1^.

For rapid chemical quenched-flow DNA unwinding experiments, the top strand (5’- GCC TCG CTG CCG TCG CCA-3’, ε_260_ = 1.53 × 10^5^ M^−1^ cm^−1^) is labeled with ^32^P at the 5’ end by T4 polynucleotide kinase (USB Corp, Cleveland, OH). The bottom strand is 5’-TGG CGA CGG CAG CGA GGC-(T)_20_-3’, ε_260_ = 3.33 × 10^5^ M^−1^ cm^−1^. A 5 μM of DNA hairpin 3’-(dT)_40_-HP10 (5’-GCC TCG CTG C-(T)_5_-G CAG CGA GGC-(T)_40_-3’, ε_260_ = 5.41 × 10^5^ M^−1^ cm^−1^) was used as a protein trap to ensure a single turn-over condition.

For single time-point studies of Rep unwinding amplitude, an 18-nucleotide ssDNA (5’-TGG CGA CGG CAG CGA GGC-3’; 5 μM, ε_260_ = 1.72 × 10^5^ M^−1^ cm^−1^) complementary to the top strand of the unwinding substrate was also added to the reaction to prevent the reannealing of the unwound product.

For fluorescence stopped-flow DNA unwinding experiments, the following DNA oligomers were used: (5’-CY5-GTT GGT CGG CAG CAG GGC-(T)_20_-3’ (ε_260,dna_ = 3.32 × 10^5^ M^−1^ cm^−1^) / 5’-GCC CTG CTG CCG ACC AAC -BHQ2 (ε_260,dna_ = 1.57 × 10^5^ M^−1^ cm^−1^)) or fluorescein, F and hexachlorofluorescein, HF (donor and acceptor) (5’-GCC TCG CTG CCG TCG CCA-Fluorescein-3’ / 5’-HF-TGG CGA CGG CAG CGA GGC-(T)_20_-3’.

For single molecule FRET experiments, the following DNA oligomers were used: 5’-Cy5-GCC CTG CTG CCG ACC AAC-3’ and 5’-Cy3-GTT GGT CGG CAG CAG GGC-(dT)_20_-3’ and their extinction coefficients are indicated above.

**Figure S1:**
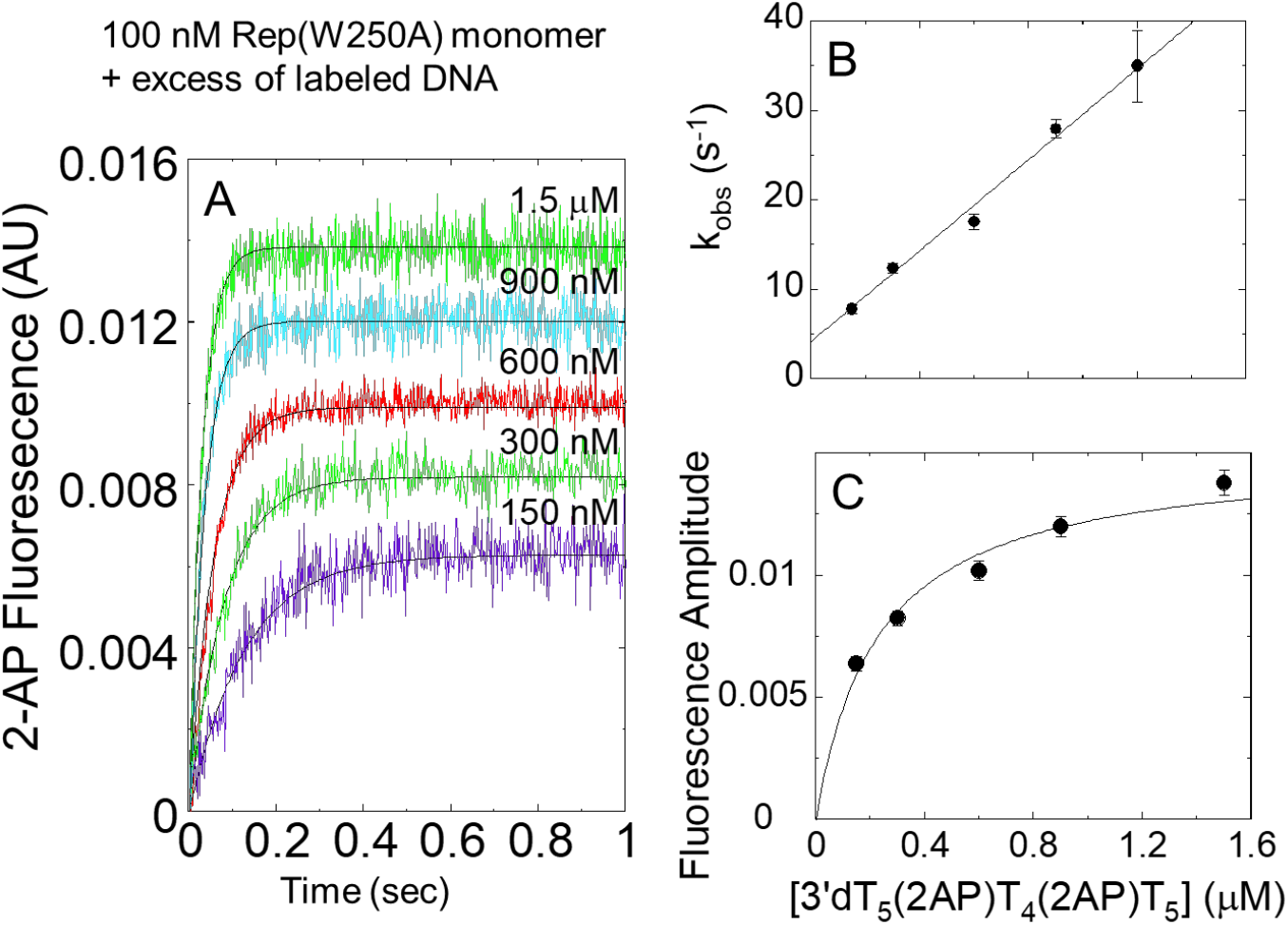
Binding kinetics of Rep(W250A) to excess 5’-(dT)_5_-(2-AP)-(dT)_4_-(2-AP)-(dT)_5_. The kinetics of Rep(W250A) binding to 5’-(dT)_5_-(2-AP)-(dT)_4_-(2-AP)-(dT)_5_ was measured by monitoring 2-AP fluorescence (λ_ex_ = 315 nm, λ_em_ ≥ 350 nm) in a buffer A 4 ºC. (**A**) 2-AP fluorescence traces upon mixing Rep(W250A) monomer (100 nM) with indicated concentrations of 5’-(dT)_5_-(2-AP)-(dT)_4_-(2-AP)-(dT)_5_. These traces have been offset to compensate for the difference in background in 2-AP fluorescence due to different DNA concentrations. The solid lines are single exponential fits to obtain the observed rate constants, k_obs_. (**B**) The observed rate constants as a function of DNA concentration. The linear fit (solid line) yields an apparent association rate constant *k*_1,app_ of 2.4±0.2 ×10^7^ M^-1^ s^-1^, and an y-intercept of 4.5±0.5 s^-1^. (**C**) The enhancement amplitude of 2-AP fluorescence as a function of DNA concentration. The data have been fitted to a square hyperbola with a zero-intercept, yielding an apparent K_d_ of 0.22±0.02 μM. The results are shown in Scheme S1

**Figure S2:**
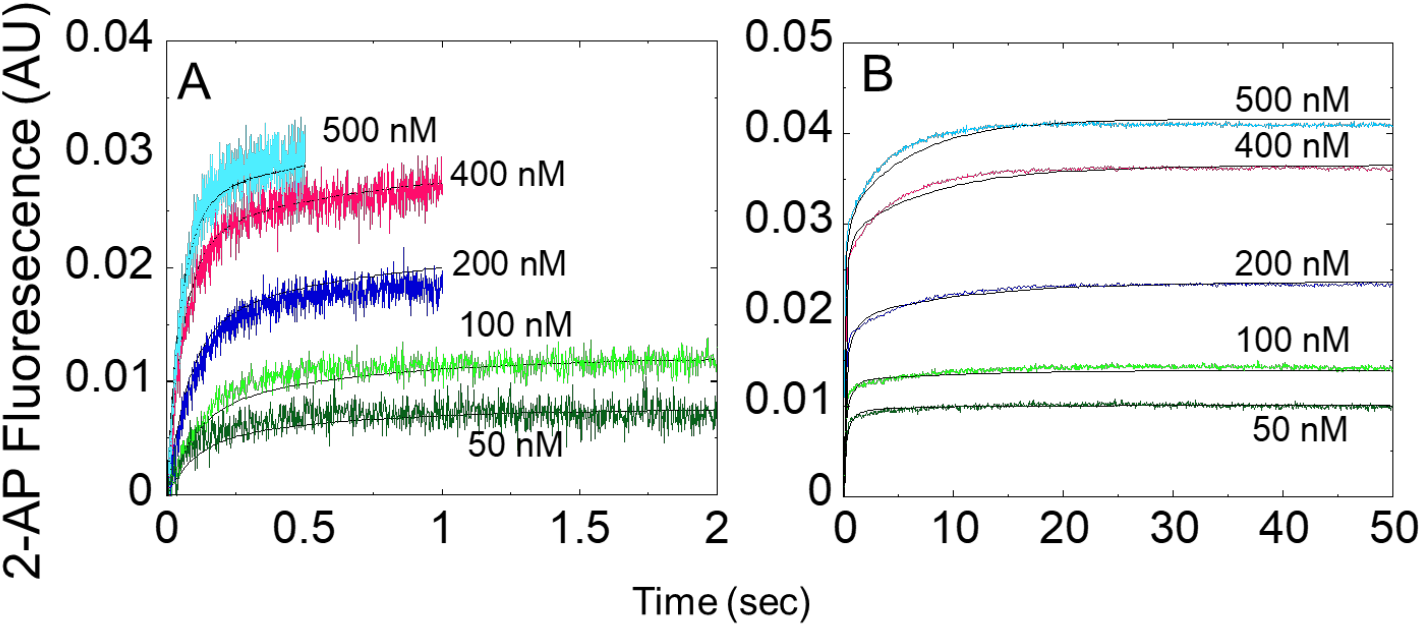
Global analysis of the kinetic fluorescence time courses of the Rep(W250A) binding to 5’-(dT)_5_-(2-AP)-(dT)_4_-(2-AP)-(dT)_5_. The stopped-flow 2-AP fluorescence traces of 100 nM 5’-(dT)_5_-(2-AP)-(dT)_4_-(2-AP)-(dT)_5_ (λ_ex_ = 315 nm, λ_em_ ≥ 350 nm) binding to various indicated concentrations of Rep(W250A) in buffer A 4 ºC. (**A**) The time courses (0 to 0.5 or 2 sec) of 2-AP fluorescence under the excess of Rep(W250A) represent the Rep(W250A) binding to DNA. (**B**) 50-sec time courses. Solid lines are simulated based on the kinetic rate constants and 2-AP fluorescence enhancement for the ssDNA-bound species obtained from global analyses. The results are summarized in the scheme S1.

**Figure S3:**
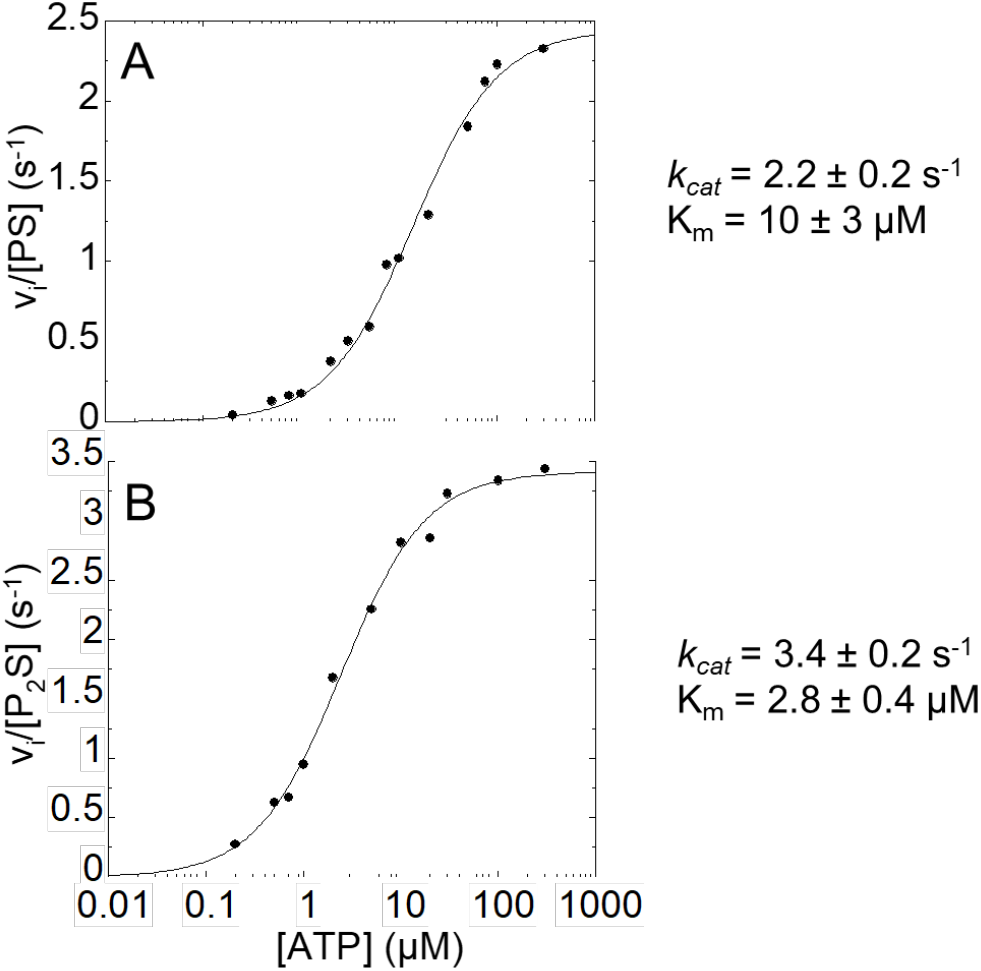
Steady-state ATPase activity of Rep(W250A) PS and P_2_S species. The experiments are conducted in buffer A 4 ºC. (**A**) Steady-state ATPase activity of Rep(W250A) PS monomer as a function of ATP concentration. The Rep(W250A) PS monomer was formed with ((dT)_16_) (4 μM) in large excess over Rep(W250A) (5 to 50 nM). Steady-state ATPase activity of Rep(W250A) P_2_S dimer as a function of ATP concentration. The Rep(W250A) P_2_S dimer was formed by mixing 800 nM Rep(W250A) with 1-10 nM ((dT)_16_). The solid lines are the fits of observed initial velocity for ATP hydrolysis, v_i_, to the Michaelis-Menten equation with *k*_cat_ = 2.2±0.2 s^-1^ and K_m_ = 10±3 μM for the Rep(W250A) PS monomer and *k*_cat_ = 3.4±0.2 s^-1^ and K_m_ = 2.8±0.4 μM for the P_2_S dimer. The W250A mutation increases the K_m_ value for PS complex while decreases the *k*_*cat*_ value for P_2_S complex as compared to the wild-type Rep.

**Figure S4:**
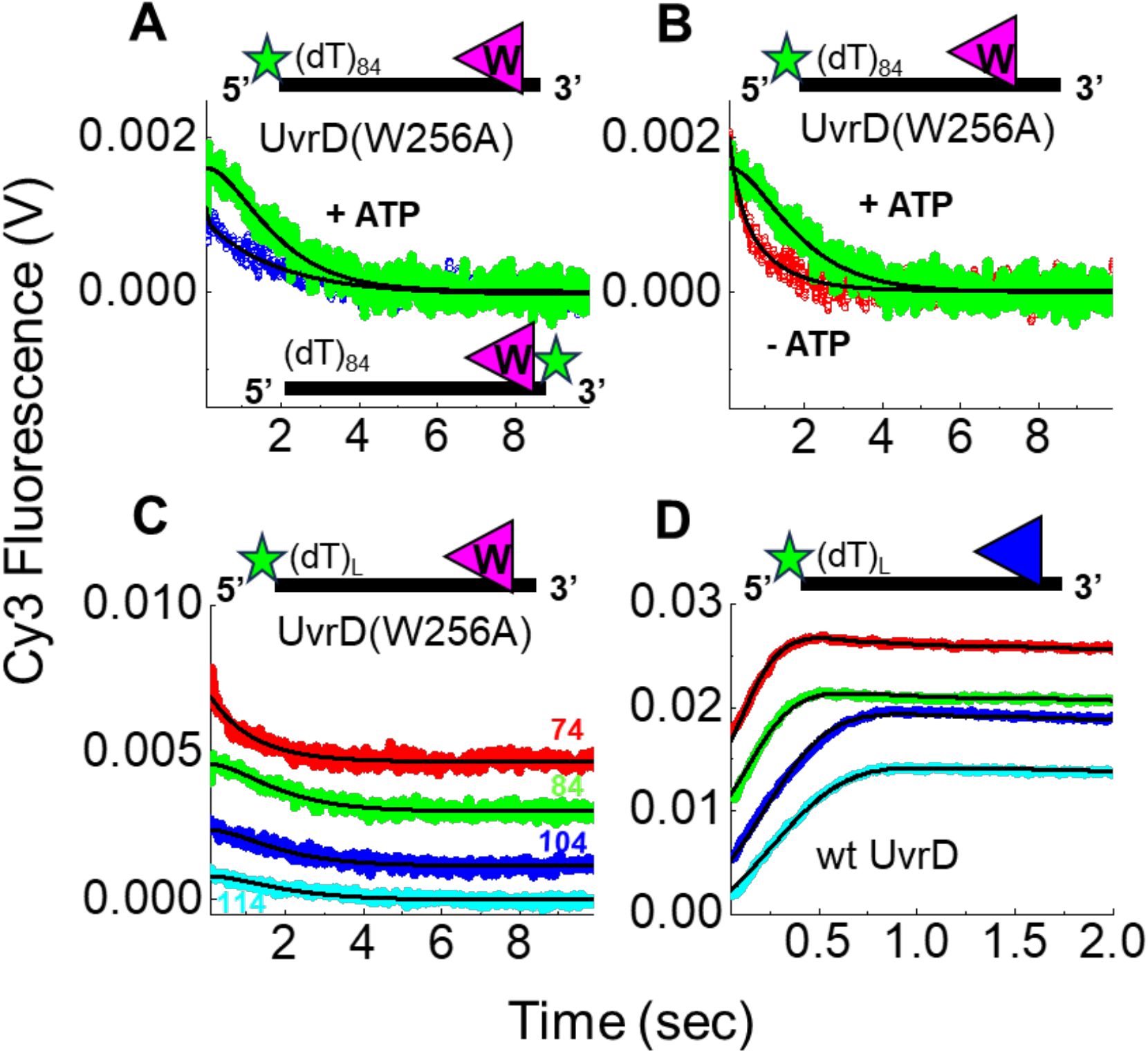
Fluorescence stopped-flow translocation of UvrD(W256A) on Cy3-labeled ssDNA. The experiments were performed in buffer U at 25 ºC by mixing a mixture (100 nM DNA + 10 nM protein) with 1.0 mM ATP, 3.0 mM MgCl_2_, 8 μM 3’-(dT)_40_-HP10 protein trap. The UvrD(W256A) translocates with a slower rate and has a lower processivity than the wtUvrD. **A**) UvrD(W256A) translocates toward the 5’ end of a ssDNA labeled with Cy3, 5’-Cy3-(dT)_84_ in the presence of 0.5 mM ATP (filled-green). When Cy3 is placed on the 3’ end ((dT)_84_-cy3-T), UvrD(W256A) translocates away from Cy3 resulting in a rapid decrease of Cy3 fluorescence (blue-open). **B**) No translocation is observed in the absence of ATP (red-open). **C**) UvrD(W256A) translocates on a 5’-Cy3-(dT)_L_ with L = 74 (red), L = 84 (green), L = 104 (blue) and L = 114 (cyan). The traces are shown with offsets for clarity. Global fits (black lines) with a two-step dissociation model (see Methods) yield a translocation rate of 7 ± 2 nts/s (Table 2). **D**) wtUvrD translocates on a 5’-Cy3-(dT)_L_ with L = 74 (red), L = 84 (green), L = 104 (blue) and L = 114 (cyan). The traces are shown with offsets for clarity. Global fits yield a translocation rate of 106 ± 2 nts/s.

**Table S1.**

| Rate Constants | Wild Type Rep<sup>#</sup> | Rep(W250A)<sup>\$</sup> |
| --- | --- | --- |
| $k_{1,app} (M^{-1}s^{-1})$ | $6.0 \pm 0.7 \times 10^7$ | $8.7 \pm 0.6 \times 10^7$ |
| $k_{-1} (s^{-1})$ | $1.2 \pm 1$ | $12 \pm 1$ |
| $k_2 + k_{-2} (s^{-1})$ | $3.9 \pm 0.6$ | $1.7 \pm 0.3$ |
| $k_3 (M^{-1}s^{-1})$ | $3.3 \pm 0.2 \times 10^5$ | $2.7 \pm 0.2 \times 10^4$ |
| $k_{-3} (s^{-1})$ | Not Determined | $0.023 \pm 0.002$ |

<sup>#</sup>Bjornson *et al.*, Biochem 1996
<sup>\$</sup>Determined in Buffer A, 4 °C
The $k_3$ of Rep(W250A) is an order of magnitude lower than that of wtRep indicating the W250 is involved in an allosteric communication between the ssDNA binding and dimerization of Rep.

**Table S2.**

| | $K_1$ | $K_2$ | $K_3$ |
| --- | --- | --- | --- |
| Rep(W250A) | $2.0 \pm 0.4 \times 10^6 M^{-1}$ | $0.8 \pm 0.04$ | $2.3 \pm 0.3 \times 10^6 M^{-1}$ |
| Wild Type Rep | $23 \pm 7 \times 10^6 M^{-1}$ | $13 \pm 6$ | $170 \pm 50 \times 10^6 M^{-1}$ |

**Table S3.**
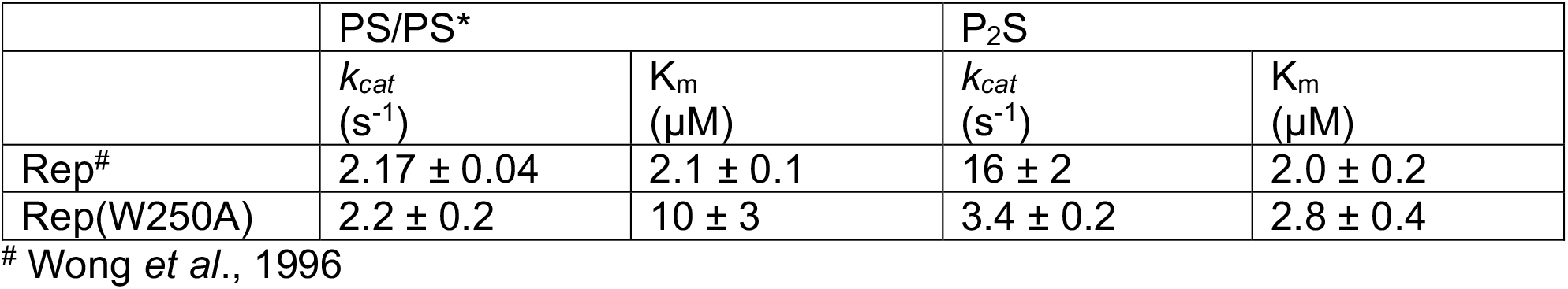
Steady-state ATPase activity of Rep(W250A) PS and P_2_S species.

|  | PS/PS<sup>*</sup> |  | P<sub>2</sub>S |  |
| --- | --- | --- | --- | --- |
| | $k_{cat}$<br>(s<sup>-1</sup>) | $K_m$<br>(μM) | $k_{cat}$<br>(s<sup>-1</sup>) | $K_m$<br>(μM) |
| Rep<sup>#</sup> | $2.17 \pm 0.04$ | $2.1 \pm 0.1$ | $16 \pm 2$ | $2.0 \pm 0.2$ |
| Rep(W250A) | $2.2 \pm 0.2$ | $10 \pm 3$ | $3.4 \pm 0.2$ | $2.8 \pm 0.4$ |

<sup>#</sup> Wong *et al.*, 1996

**Table S4:** Summary of smFRET DNA unwinding experiments for Rep homo and heterodimers. There are no systematic variations of the percentages of molecules unwound for homodimers and heterodimers at 3 ATP concentrations (18-28%). Biotinylated-Rep (bioRep) is immobilized on the surface.

|  | bioRep + Rep |  |  | bioRep + Rep(W250A) |  |  | bioRep(W250A) + Rep |  |  |
| --- | --- | --- | --- | --- | --- | --- | --- | --- | --- |
| ATP (μM) | 5 | 10 | 50 | 5 | 10 | 50 | 5 | 10 | 50 |
| Total Number of Molecules | 114 | 104 | 170 | 128 | 98 | 103 | 95 | 115 | 127 |
| Number of molecules unwound | 21 | 22 | 34 | 22 | 25 | 22 | 26 | 23 | 28 |
| Percent unwound | 18.4 | 21.1 | 20.0 | 17.1 | 25.5 | 21.4 | 27.4 | 20.0 | 22.0 |

**Table S5:** Summary of smFRET DNA unwinding experiments for UvrD homo and heterodimers. The percentages of molecules unwound are slightly lower for the heterodimers at higher ATP concentrations. Biotinylated-UvrD (bioUvrD) is immobilized on the surface.

|  | bioUvrD + UvrD |  |  | bioUvrD + UvrD(W256A) |  |  |
| --- | --- | --- | --- | --- | --- | --- |
| ATP (μM) | 5 | 10 | 25 | 5 | 10 | 25 |
| Total Number of Molecules | 43 | 87 | 67 | 72 | 74 | 159 |
| Number of molecules unwound | 14 | 30 | 20 | 24 | 14 | 25 |
| Percent unwound | 32.6 | 34.5 | 29.9 | 33.3 | 18.9 | 15.7 |

#### Scheme S1

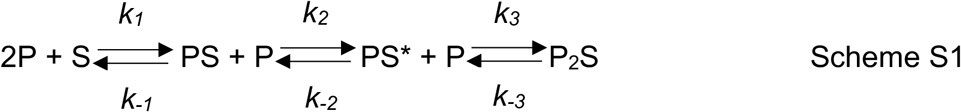

Where P is the Rep monomer, and S is the DNA. Under pseudo-first order conditions, the observed rate constants for the three phases can be approximated by Equations i, ii, and iii.

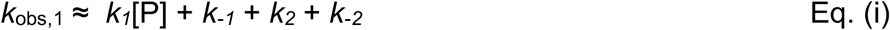

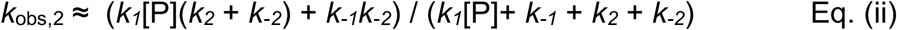

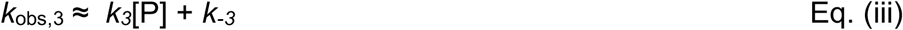

At high protein [P] concentration, *K*_obs_,_2_ ≈ *k*_2_ + *k*-_2_ I .unit Communica

Kinetic constants for ssDNA binding and induced dimerization of Rep or Rep(W250A). Rep(W250A) binds weaker to ssDNA than wt Rep.

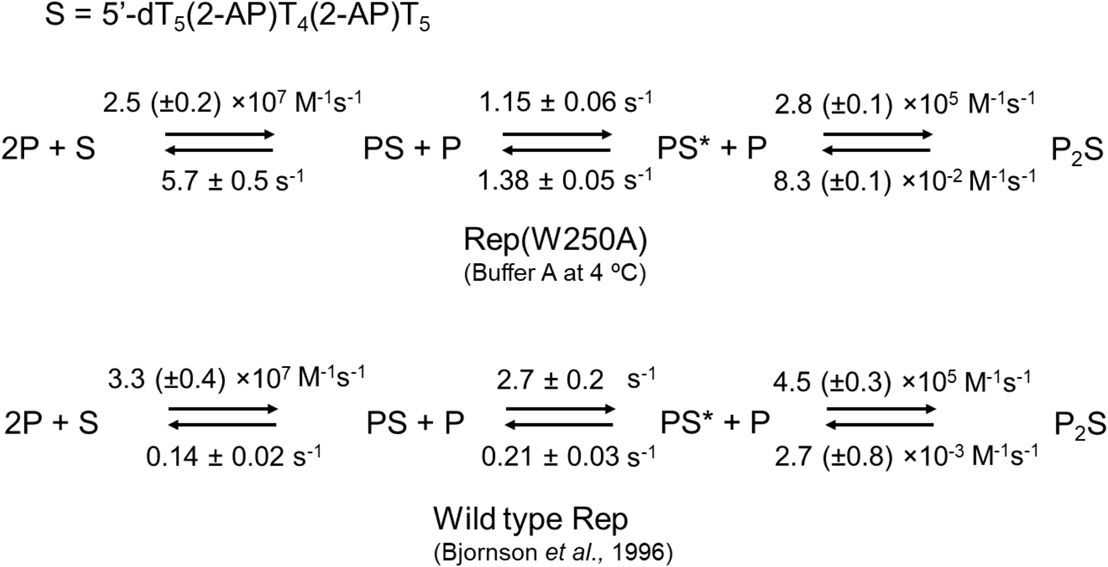

